# F-Actin Bending Facilitates Net Actomyosin Contraction By Inhibiting Expansion With Plus-End-Located Myosin Motors

**DOI:** 10.1101/2021.10.03.462946

**Authors:** Alexander K. Y. Tam, Alex Mogilner, Dietmar B. Oelz

**Affiliations:** School of Mathematics and Physics, The University of Queensland, St Lucia, Queensland 4072, Australia; Courant Institute of Mathematical Sciences, New York University, New York, NY, USA; School of Mathematical Sciences, Queensland University of Technology, Brisbane, Queensland 4000, Australia

**Keywords:** actomyosin, curve-straightening flow, energy functional, gradient flow, stress tensor, asymptotic analysis

## Abstract

Contraction of actomyosin networks underpins important cellular processes including motility and division. The mechanical origin of actomyosin contraction is not fully-understood. We investigate whether contraction arises on the scale of individual filaments, without needing to invoke network-scale interactions. We derive discrete force-balance and continuum partial differential equations for two symmetric, semi-flexible actin filaments with an attached myosin motor. Assuming the system exists within a homogeneous background material, our method enables computation of the stress tensor, providing a measure of contractility. After deriving the model, we use a combination of asymptotic analysis and numerical solutions to show how F-actin bending facilitates contraction on the scale of two filaments. Rigid filaments exhibit polarity-reversal symmetry as the motor travels from the minus to plus-ends, such that contractile and expansive components cancel. Filament bending induces a geometric asymmetry that brings the filaments closer to parallel as a myosin motor approaches their plus-ends, decreasing the effective spring force opposing motor motion. The reduced spring force enables the motor to move faster close to filament plus-ends, which reduces expansive stress and gives rise to net contraction. Bending-induced geometric asymmetry provides both new understanding of actomyosin contraction mechanics, and a hypothesis that can be tested in experiments.

## 1 Introduction

The mechanics of actin filaments and myosin motor proteins in the cell cortex underpins movement [1] and division [2] of biological cells. Early breakthroughs in understanding actomyosin dynamics occurred in muscle cells [3], in which actin and myosin form sarcomere structures. Sarcomeres involve filaments aligned in parallel with minus-ends in the centre and plus-ends pointing outwards. Relative motion of myosin motors towards filament plus-ends subsequently generates contraction by pulling filaments inwards. This mechanism is known as sliding filament theory [4]. However, actomyosin networks in the cell cortex are disordered, with filaments distributed at random. Experiments [5, 6] and simulations [7, 8] have shown that disordered actomyosin networks also contract [9]. According to sliding filament theory, filament pairs produce expansion if myosin motor proteins localise close to plus-ends, or contraction if myosin motor proteins localise close to minus-ends. In disordered actomyosin networks, motors localise near plus-ends and minus-ends with equal probability. Therefore, sliding filament theory alone cannot explain disordered network contraction. The origin of contraction in disordered actomyosin networks remains an active field of research.

Filament bending flexibility is commonly-hypothesised as a source of asymmetry that might explain contraction of disordered actomyosin networks [7, 10–13]. Actin filaments are semi-flexible [14, 15], such that they undergo small but significant bending [10, 16]. Filament semi-flexibility is irrelevant in sarcomeres with parallel arrays of straight filaments, but is important for disordered networks in which motors can cross-link filaments at arbitrary angles and generate torque. Previous experimental and theoretical studies in disordered networks suggest that filament buckling gives rise to contraction. These studies show that filaments can sustain longitudinal tension, but buckle under longitudinal compression [10, 12, 15, 17–24]. This buckling mechanism can generate network-scale bias to contraction over expansion [15]. Other studies have considered a related filament bending mechanism [7, 13, 25–28] as a source of force asymmetry. Bending involves applying forces that pluck filaments transversely, in contrast to the longitudinal forces involved with buckling. Lenz [25] showed that filament bending produces forces that exceed those involved with longitudinal buckling, and Tam, Mogilner, and Oelz [7] showed that this mechanism facilitates network-scale contraction. A pertinent question is whether the force asymmetry provided by bending or buckling applies at the microscopic scale [11, 25, 29], or whether long-range effects transmit contractile force through the network, without requiring a microscopic asymmetry [22].

One approach to understand microscopic filament dynamics is to model a single filament as an inextensible elastic rod, as in ‘worm-like chain’ models [16, 25]. Broedersz and Mackintosh [16] used this approach to identify an asymmetry under extension and compression. Other authors have considered structures consisting of two-filaments and an attached motor [15, 25, 29, 30]. Lenz [25] reported that disordered networks of rigid filaments with polarity-reversal symmetry (*i*.*e*. any configuration of filaments is equally likely as the same configuration with minus and plus-ends reversed) generate zero net contraction. Lenz [25] also showed that filament bending gives rise to contraction in a two-filament system, and is the dominant mechanism of contraction for experimentally-feasible parameters. During motor-induced deformation, semi-flexible actin filaments evolve to minimise their bending energy. Therefore, we model semi-flexible filament evolution as a curve-straightening flow. Mathematically, curve-straightening refers to deformation of curves in ℝ^2^ by decreasing their total squared curvature. Curve-straightening problems have been investigated since the 1980s [31–34]. Wen [35, 36] used the indicatrix representation and *L*^2^-gradient flow of the squared curvature functional to derive a fourth-order, semilinear parabolic partial differential equation (PDE) for the evolution of the curve. Oelz [37] extended this work to model an open curve. However, theoretical analysis of curve-straightening flows is mostly limited to single curves, rather than the two-curve representations necessary to model a pair of filaments.

We extend previous curve-straightening models to derive a coupled PDE system for two symmetric, semi-flexible filaments with a myosin motor attached at their intersection. After obtaining the governing equations, we describe how to obtain the stress tensor, assuming the two filaments reside in a homogeneous background material. We then use asymptotic analysis and numerical solutions to provide a detailed explanation of how filament bending facilitates contraction on the two-filament scale. Our analysis suggests a contraction mechanism based neither on filament buckling, nor intrinsic force asymmetry where bending generates contraction. Instead, filament semi-flexibility creates a geometric asymmetry that inhibits expansion. Rigid filaments exhibit polarity-reversal symmetry, whereby contraction associated with a minus-end-located motor balances with expansion associated with a plus-end-located motor. Allowing filaments to bend breaks this symmetry, and the filaments become closer to parallel as the motor approaches the plus-ends. This decreases the spring force through the motor, enabling the motor to move faster close to the plus-ends. Fast motor motion inhibits expansive stress, and gives rise to net contraction. Our analysis provides a new hypothesis for bending-induced actomyosin contraction, and shows how contraction can occur on the microscopic two-filament scale.

## 2 Mathematical Model

We develop a mathematical model for a myosin motor attached to two overlapping actin filaments. We represent filaments as open curves in ℝ^2^, and denote their positions by *z*_*i*_(*s*(*t*), *t*) = (*x*_*i*_(*s*(*t*), *t*), *y*_*i*_(*s*(*t*), *t*)), for *i* = 1, 2 (see Figure 2.1). The variable *t* denotes time, and *s* ∈ [0, *L*_*i*_] is the arc length parameter, where *L*_*i*_ is the length of the *i*-th filament. Actin filaments are polarised, so we adopt the convention that *s* = 0 corresponds to the filament minus-end, and *s* = *L*_*i*_ corresponds to the plus-end. Since non-muscle myosin thick filaments are short compared to actin filaments [38], we represent the myosin motor as a point object existing at the intersection between the two filaments. We track its position by introducing the variables *m*_*i*_(*t*) ∈ [0, *L*_*i*_], such that *s* = *m*_*i*_ is the position of the motor head attached to the *i*-th filament. We assume that no other proteins, for example cross-linkers, are present.

**Figure 2.1:**
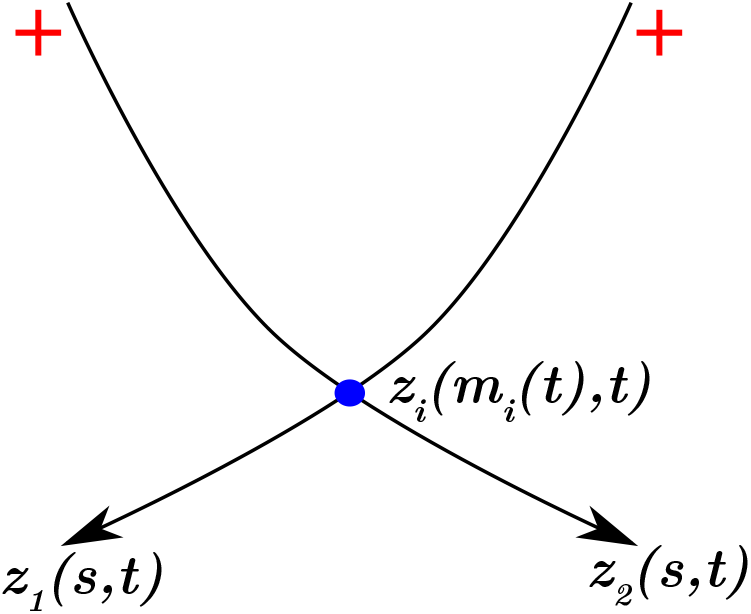
Schematic representation of two actin filaments with a myosin motor attached at their intersection. Filaments are the curves *z*_1_ and *z*_2_, and arrow heads represent minus-ends. The myosin motor protein is represented by the blue dot.

### 2.1 Discrete Force-Balance Equations

We express the mathematical model for the filament and motor mechanics as a system of force-balance equations,

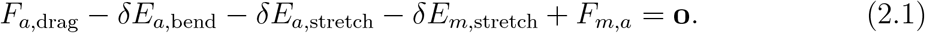

The first three terms in (2.1) describe the drag, bending, and longitudinal stretching forces respectively on actin filaments. The fourth term represents longitudinal stretching along the myosin motor, and the final term describes forces between filaments and motors. We represent bending and stretching forces as the variation of potential energy, where terms involving *δ* denote variations. We formulate the force-balance equations (2.1) as a minimisation problem. This involves constructing a time-discrete scalar functional that contains a contribution for each force term in (2.1),

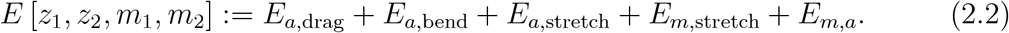

In (2.2), *E*_*a*,bend_, *E*_*a*,stretch_, and *E*_*m*,stretch_ are the potential energies associated with filament bending, filament stretching, and motor stretching respectively. The terms *E*_*a*,drag_ and *E*_*m,a*_ are pseudo-energy terms with variations that correspond to finite-difference approximations of *F*_*a*,drag_ and *F*_*m,a*_, which cannot be interpreted as variations of potential energy.

The first term, *E*_*a*,drag_, describes drag friction between filaments and a passive background medium. Drag acts uniformly along the filaments and opposes filament motion. The term to represent drag between filaments and the background medium is

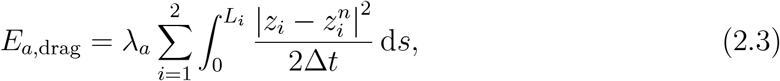

where *λ*_*a*_ is the filament drag coefficient, Δ*t* is the time step size, and the superscript *n* refers to the previous time step in the discrete formulation, 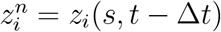. We model bending of semi-flexible actin filaments via the elastic potential energy

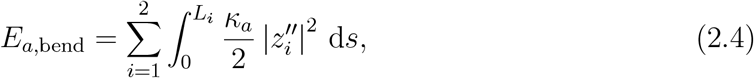

where *κ*_*a*_ is the flexural rigidity, and primes denote differentiation with respect to arc-length, *s*. We assume that *κ*_*a*_ is constant, and the same for both filaments. We obtain a term for filament stretching by assuming that actin filaments are inextensible. To model this, we ensure that 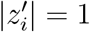 at every point along the filaments using the penalisation term

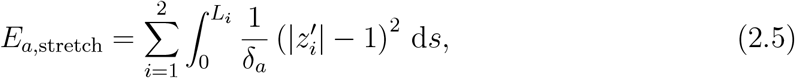

where *δ*_*a*_ is an arbitrarily small parameter that enforces the inextensibility constraints.

The remaining two terms in (2.2) describe how motors contribute to the mechanics. To model motor stretching, we introduce another penalising potential,

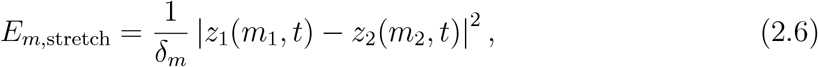

where *δ*_*m*_ is an arbitrarily small parameter that penalises deviation from the constraint *z*_1_(*m*_1_, *t*) = *z*_2_(*m*_2_, *t*) stating that motors are to remain point objects. The final term in (2.2) describes interactions between filaments and motors. We assume that motors obey a linear (affine) force–velocity relation [39], illustrated in Figure 2.2. This law integrates the multitude of force contributions exerted by myosin heads which decorate the myosin thick filament and interact with actin filaments [40]. Subject to zero force (when the motor is unextended), motors move with speed *V*_*m*_. As the motor extends, the spring force through the motor increases. We assume that motor velocity varies linearly with the spring force through the motor. If the force through the motor exceeds *F*_*s*_, the stall force, then the motor velocity is zero. The corresponding pseudo-energy term consists of a linear term, and a quadratic drag-like term for the velocity reduction caused by the force through the motor,

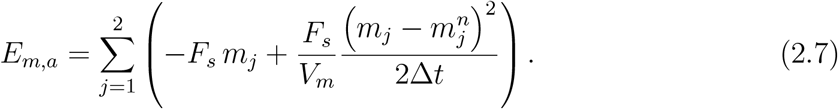

**Figure 2.2:**
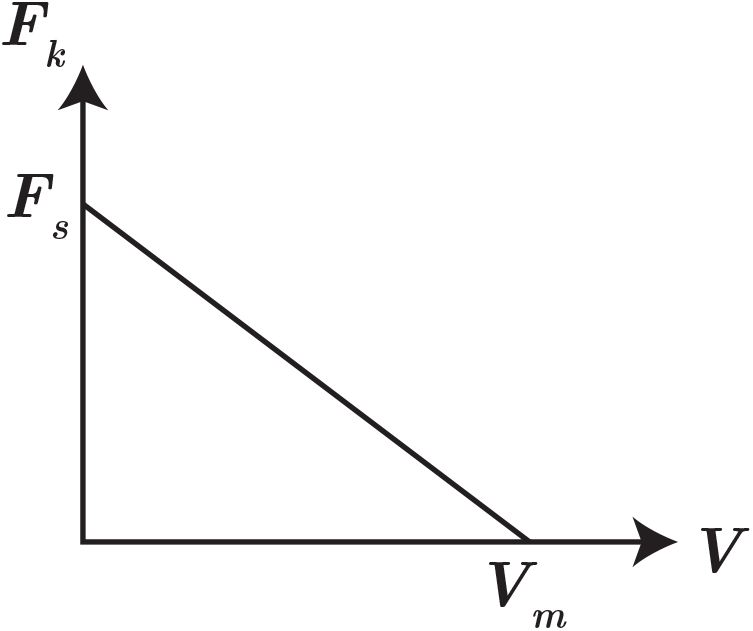
The linear force–velocity relationship for myosin motors bound to actin filaments. An unextended motor subject to zero force moves towards filament plus-ends with the free-moving velocity, *V*_*m*_. If the spring force through the motor exceeds the stall force *F*_*s*_, the motor does not move.

With the definition of (2.3)–(2.7), minimising the functional (2.2) for fixed Δ*t* provides a time-implicit numerical method to solve the force-balance equations (2.1) for the filament and motor positions.

### 2.2 Governing Partial Differential Equations

Formulating the model as a minimisation problem enables us to derive a continuum model for the filament and motor positions based on the discrete formulation in §2.1. The derivation is based on the following variational principle. Given known data 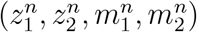 at the discrete point in time *n*, the solution at the following point in time minimises the functional (2.2), that is

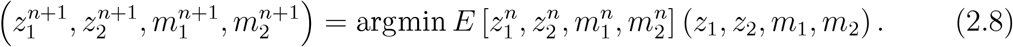

We obtain force-balance equations by setting to zero the functional derivatives of (2.2) with respect to filament and motor positions. Subsequently, we write

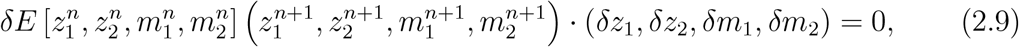

where terms involving *δ* denote the variation of the respective quantity. Minimising the functional (2.2) enables us to write the force-balance equations (2.1) in terms of *z*_1_, *z*_2_, *m*_1_, and *m*_2_. We obtain the governing equations by evaluating (2.9) and matching coefficients of *δz*_1_, *δz*_2_, *δm*_1_, and *δm*_2_. On taking the formal continuum limit Δ*t* → 0, for which 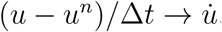, we obtain the system of PDEs

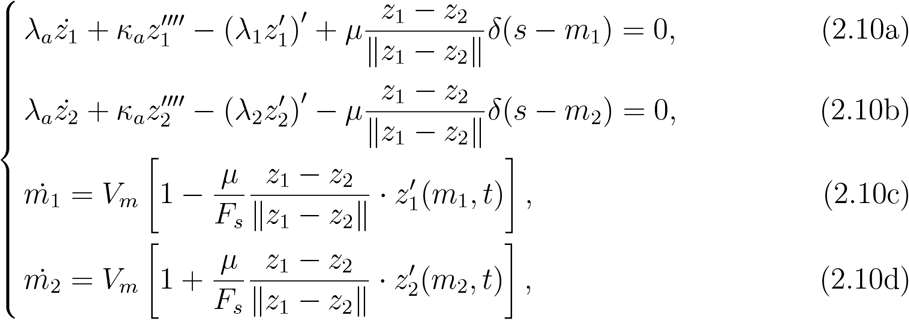

where primes denote differentiation with respect to arc length, dots represent derivatives with respect to time, and *δ*(·) is the Dirac delta function (not to be confused with variation). Equations (2.10) are a system of continuum force-balance equations for the filament and motor positions. They are formulated in a formal limit where *δ*_*a*_ and *δ*_*m*_ are small, and the force coefficients 1*/δ*_*a*_ and 1*/δ*_*m*_ in the variations of the penalising potentials (2.5) and (2.6) are replaced by the Lagrange multipliers *λ*_1_, *λ*_2_, and *μ*. Note that the sign of *z*_1_ − *z*_2_ in (2.10) will be absorbed by *μ*. As a consequence, solutions satisfy the constraints

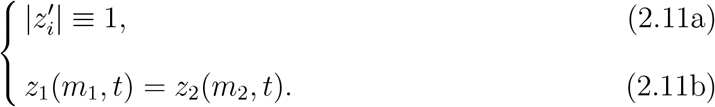

The equations are subject to the boundary and initial conditions

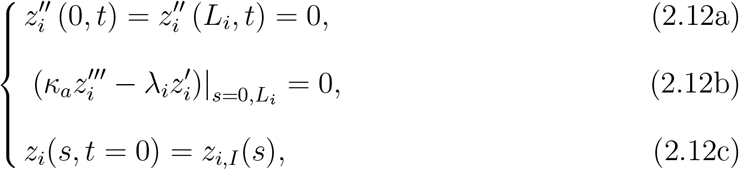

where the subscript *I* represents an initial quantity. A detailed derivation of (2.10) and (2.12) is provided in Appendix A.

### 2.3 Calculation of Forces and Stress

An objective of this work is to describe how the filament and motor motion governed by our model generates contractile and expansive forces. We assume the two-filament–motor structure is immersed in a dense network of cross-linked filaments covering a rectangular domain. A pair of actin filaments can locally deform the background network in which it is immersed. However, we assume the background can only undergo uniform elongation and shearing. A scenario supporting this assumption is that the background network consists of numerous two-filament–motor assemblies, all with the same shape as the reference pair that we describe explicitly (Figure 2.3B). In this scenario, deformations occur equally everywhere in the domain, and the background network remains homogeneous. If the background is homogeneous, we can associate the tension at the boundaries of the domain with the stress being generated by the reference pair of actin filaments (Figure 2.3A).

**Figure 2.3:**
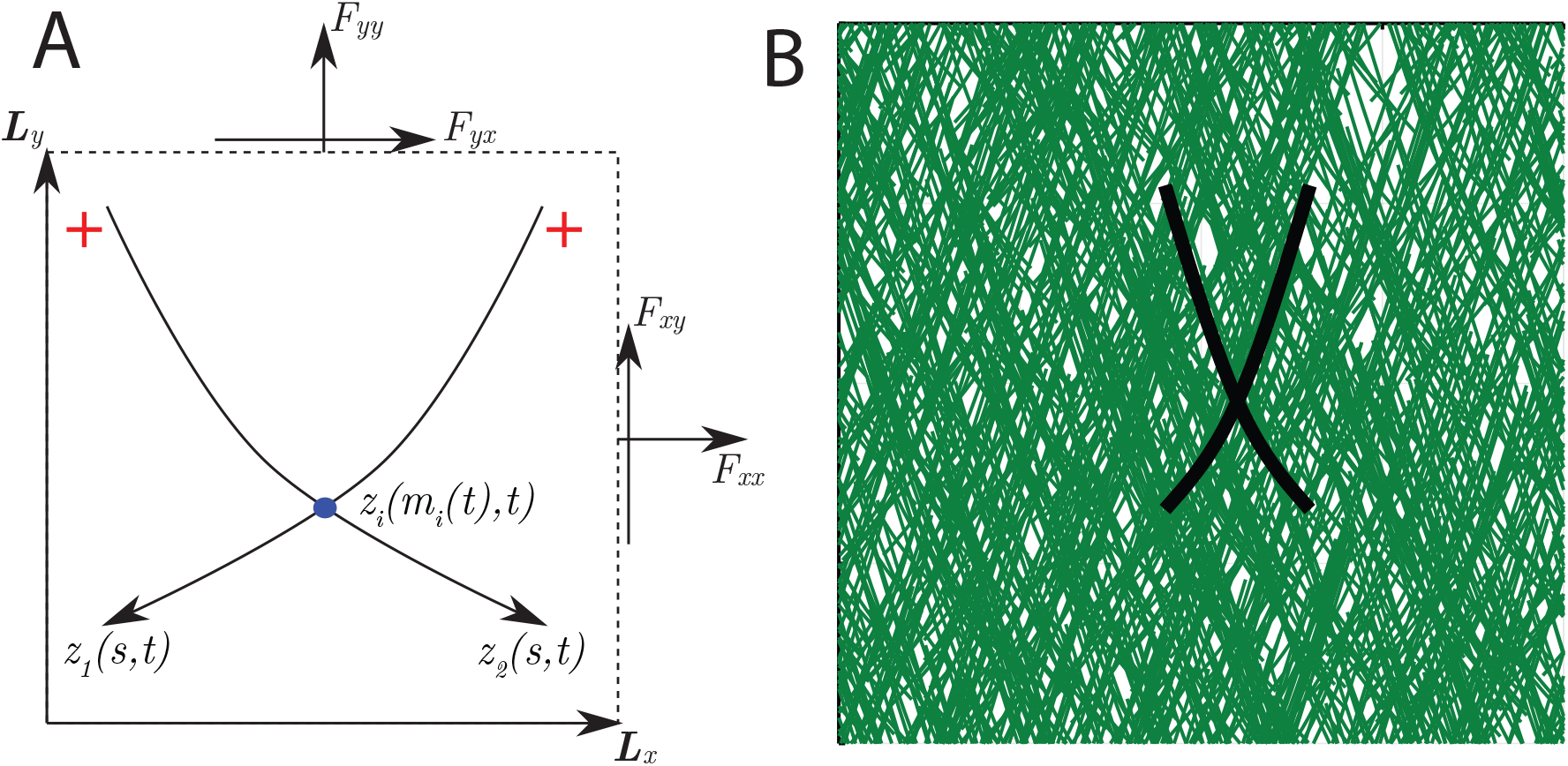
Schematic representation of a two-filament–motor system existing within a rectangular homogeneous background medium. A: The filaments are the curves *z*_1_ and *z*_2_, and arrow heads represent filament minus (pointed) ends. Myosin motor proteins are represented by blue dots, and initially appear at the intersection between the two filaments. B: Visualisation of the two-filament–motor structure embedded in a dense background network consisting of numerous assemblies (green lines) with the same shape as the reference pair (black lines).

To quantify contraction, we compute the stress tensor for a small rectangular region of background material that encloses the two-filament–motor structure. The adjacent sides of the rectangle are given by the vectors ***L***_*x*_ = (*L*_*xx*_, *L*_*xy*_)^*T*^, and ***L***_*y*_ = (*L*_*yx*_, *L*_*yy*_) as shown in Figure 2.3A. We compute the vectors ***F***_*x*_ = (*F*_*xx*_, *F*_*xy*_) and ***F***_*y*_ = (*F*_*yx*_, *F*_*yy*_), also shown in Figure 2.3A. These vectors are the force components acting on the domain boundaries that must be applied to prevent uniform elongation and shear deformations. These forces sum the contributions of both filaments and the motor, and provide a measure of net contractility. To derive closed form expressions for these forces, we return to the discrete formulation. We obtain ***F***_*x*_ and ***F***_*y*_ by first adding extra terms to the functional (2.2), and defining

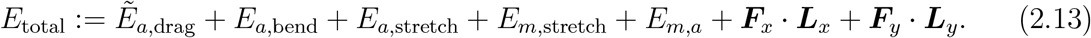

In (2.13), we use the modified drag term

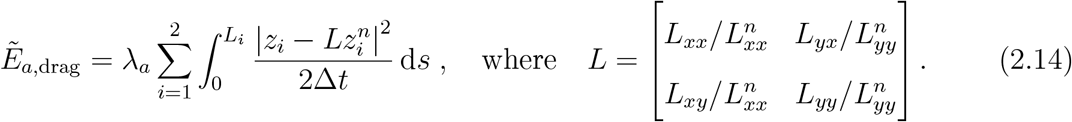

The matrix *L* represents the transition from the coordinate frame 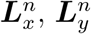 at time *n*, to the new coordinate frame ***L***_*x*_, ***L***_*y*_, corresponding to a rectangle that has undergone uniform shearing and elongation. If we impose 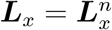 and 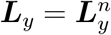, the vectors 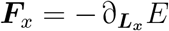 and 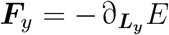 represent Lagrange multipliers that enforce the constant domain size and shape constraints. We assume that any possible deformations of the rectangle are small, such that Cauchy stress theory applies. The two-dimensional state of stress in the domain is then given by the stress tensor,

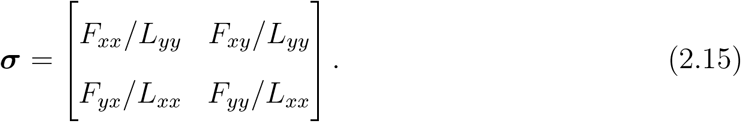

The bulk stress,

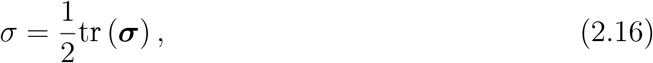

then provides a measure of the contraction or expansion generated by the two-filament– motor system. By convention, negative *σ* indicates contraction, and positive *σ* indicates expansion. The quantity *σ* is invariant to domain rotations, and equal to the average of the eigenvalues of ***σ***. The associated eigenvectors of ***σ*** are the principal stress directions, which indicate the directions of maximum contraction or expansion.

To obtain an explicit expression for the bulk stress, *σ*, in terms of the filament positions, we differentiate the functional (2.13) with respect to ***L***_*x*_ and ***L***_*y*_. This yields

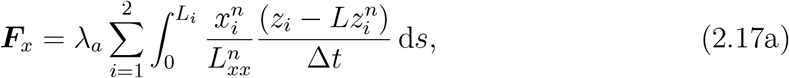

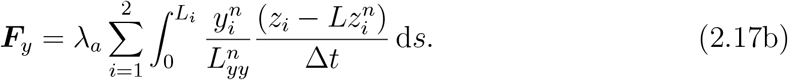

Applying the formal continuum limit Δ*t* → 0, *L* → ***I***, and 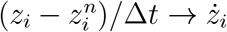, we obtain

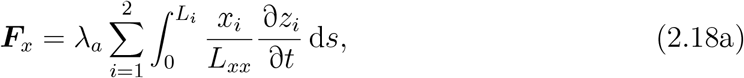

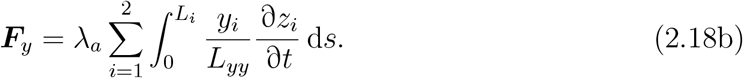

Evaluating the bulk stress (2.16) then yields

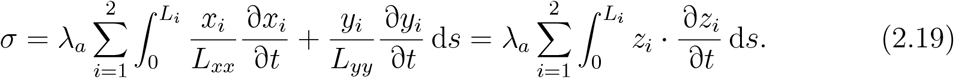

Furthermore, the expressions (2.18) confirm that

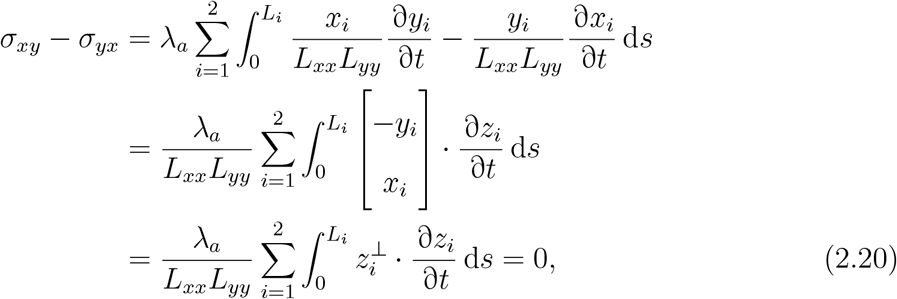

where 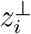 denotes a vector orthogonal to *z*_*i*_, and we obtain the result by substituting (2.10) for 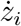. The stress tensor (2.15) is thus symmetric, and the bulk stress *σ* is equal to the average of the eigenvalues of ***σ***.

### 2.4 Nondimensionalisation

We nondimensionalise the PDE model (2.10)–(2.12) by introducing the length and time scales

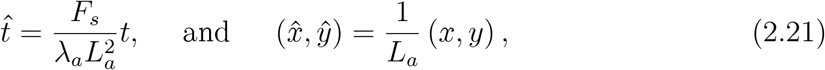

where hats represent dimensionless variables, and *L*_*a*_ is a characteristic filament length. The dimensionless model is then (dropping hats for convenience)

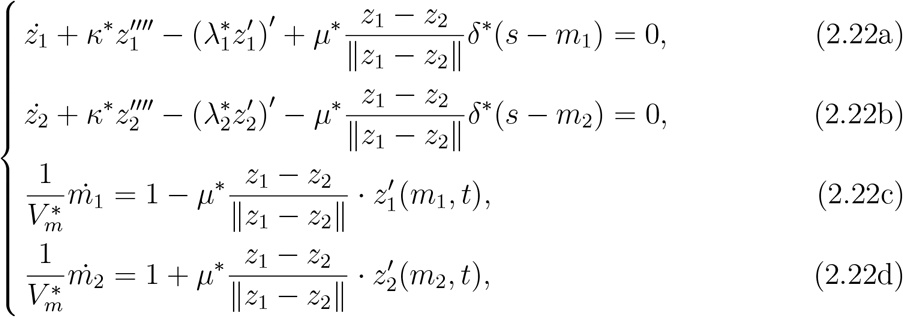

subject to the boundary and initial conditions

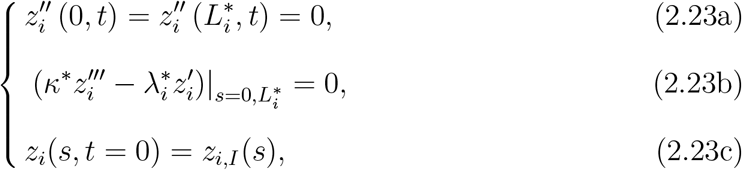

and the constraints

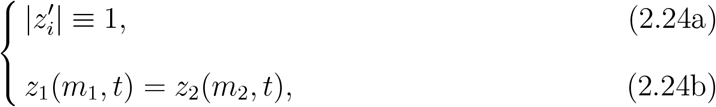

where 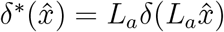 is a scaled Dirac delta function. The dimensionless parameters and forces are

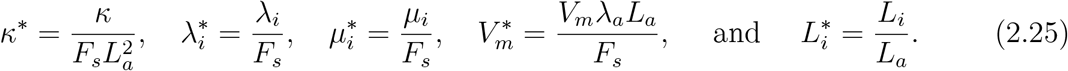

### 2.5 Model Simplification

Before obtaining asymptotic and numerical results, we consider a simplification to the dimensionless model (2.22)–(2.24). First, we assume that the two filaments are symmetric about the vertical, that is

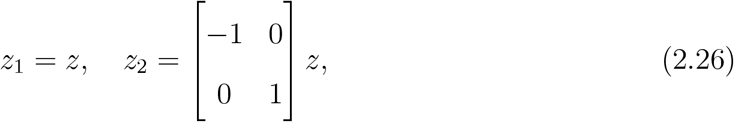

and have identical length *L*_*a*_ = *L*_1_ = *L*_2_. Symmetry also implies that the relative position of the motor is the same for both filaments, *m*_1_ = *m*_2_ = *m*, and that 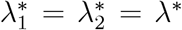. To simplify the motor dynamics (2.22c) and (2.22d), we impose *V*_*m*_ → ∞. Finally, we rewrite dimensionless flexural rigidity according to *κ*\* = 1*/ε*, indicating that the flexural rigidity is large (*ε* ≪ 1), and that the filaments undergo only minor bending. On applying these simplifications, the dimensionless model (2.22) becomes (dropping asterisks on dimensionless parameters)

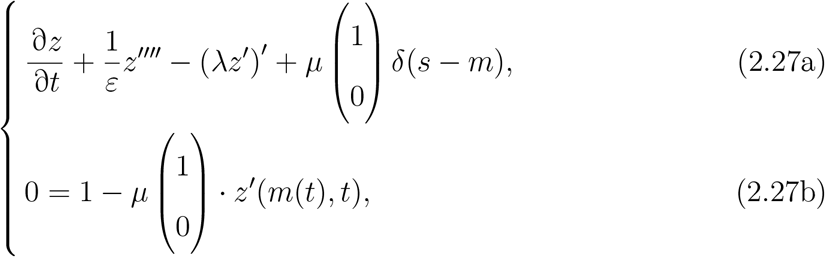

subject to the boundary and initial conditions

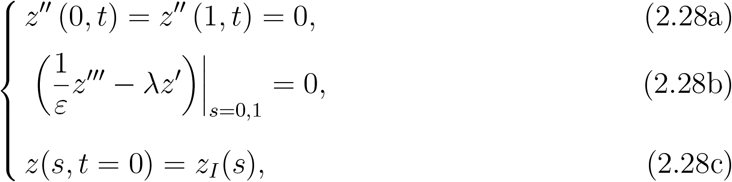

and the constraints

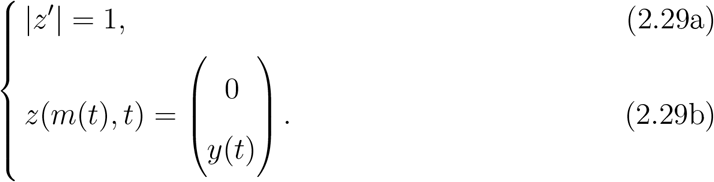

Equations (2.27a) and (2.27b) describe the filament and motor evolution respectively, under the simplifying assumptions. Equations (2.27a) and (2.27b), with the boundary and initial conditions (2.28), and constraints (2.29), complete our simplified model.

The dimensionless bulk stress (2.19) for the simplified model becomes

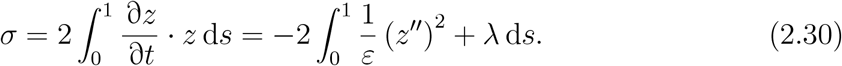

To obtain a measure of net stress, we integrate *σ* over the time between motor attachment and detachment. This yields

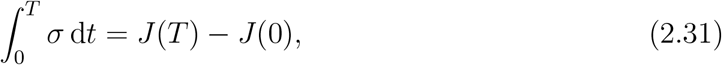

where

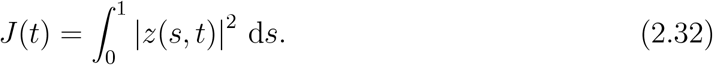

The quantity *J* (*T*) − *J* (0) describes the net, time-aggregated stress that the two filaments produce between motor attachment and detachment. This quantity will be important in our asymptotic and numerical investigation on how filament bending affects contraction.

## 3 Results and Discussion

We analyse the simplified model derived in §2.5 to quantify how filament flexibility gives rise to contractile stress. First, we use asymptotic analysis to obtain an explicit approximation to the solution in the limit of infinite flexural rigidity. Through the leading-order problem, we show that a rigid two-filament–motor structure with polarity-reversal symmetry produces zero net stress. The first-order problem gives rise to a system of differential equations that governs the dynamics with small filament bending. Second, we obtain numerical solutions to validate the asymptotic analysis, and investigate solutions beyond the large flexural rigidity limit. These solutions reveal that contraction arises from a geometric asymmetry, whereby filaments become more parallel as the motor approaches the plus-ends. This inhibits expansion associated with plus-end-located myosin motors. Since contraction associated with minus-end-located motors is unaffected, the net outcome is a contractile two-filament–motor structure.

### 3.1 Asymptotic Analysis

We construct an asymptotic approximation to the solution of the simplified symmetric model (2.27)–(2.29). Asymptotic analysis involves expanding variables in powers of *ε*,

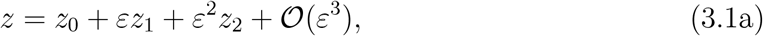

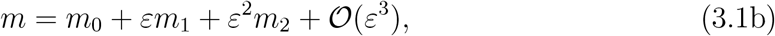

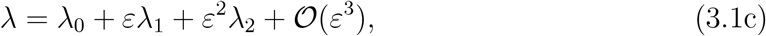

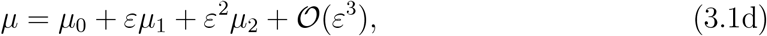

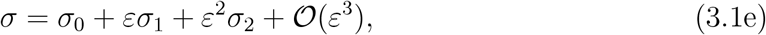

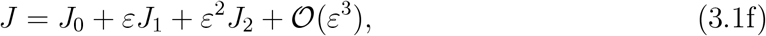

as *ε* → 0. On substituting the asymptotic series (3.1) into the model (2.27)–(2.29), the leading-order solution is the evolution of two rigid filaments with infinite resistance to bending. The first-order problem describes how small, non-zero bending affects the dynamics and stress. We present the key results and arguments in subsequent subsections, and give full details of the computations in Appendix B.

#### 3.1.1 Leading-Order Solution

The leading-order solution describes the evolution of rigid filaments. To solve for *z*_0_, we consider the balance at 𝒪(1*/ε*). This yields

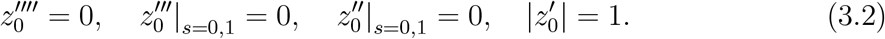

The solution to the 𝒪(1*/ε*) problem (3.2) is a straight filament, whose direction we parameterise by the filament angle, *θ/*2, measured from the positive vertical axis (see Figure 3.1). We write

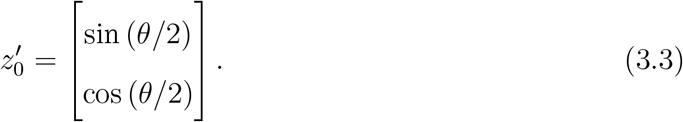

**Figure 3.1:**
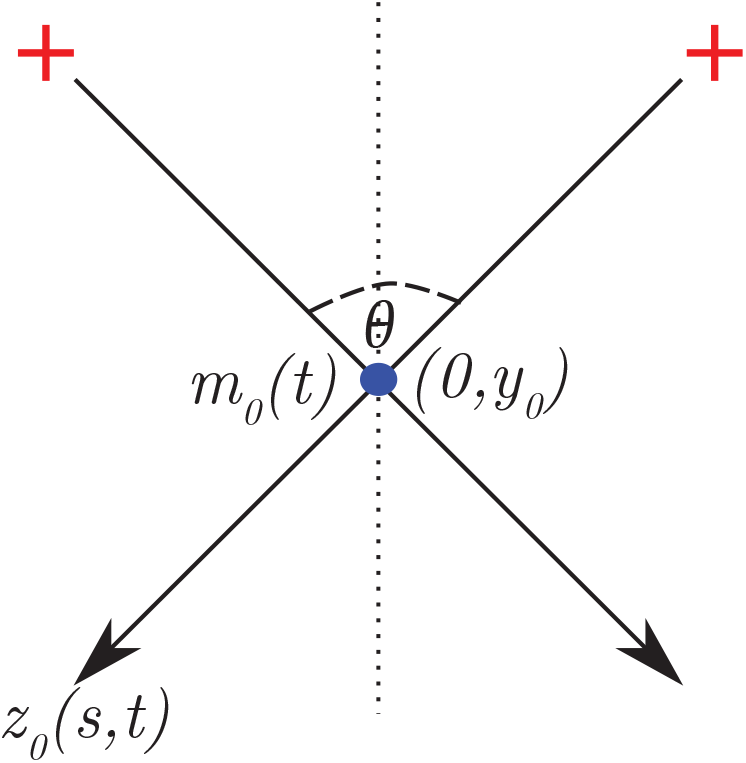
Schematic of a two-filament–motor system with rigid symmetric actin filaments. The myosin motor has relative position *m*_0_(*t*), and physical position (0, *y*_0_(*t*)). Filaments are symmetric about the dashed vertical line, which is the positive *y*-axis. The angle between the filaments is *θ*, such that the angle between a filament and the *y*-axis is *θ/*2.

For a filament orthogonal to *z*_0_ we use the notation

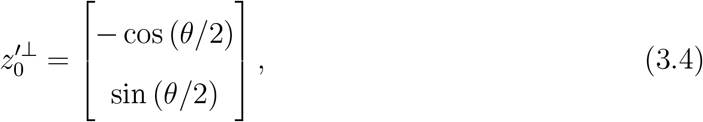

where the symbol ^⊥^ denotes rotation to the left by *π/*2. A suitable ansatz for the position of a rigid filament solution satisfying the constraint (2.29b) is then

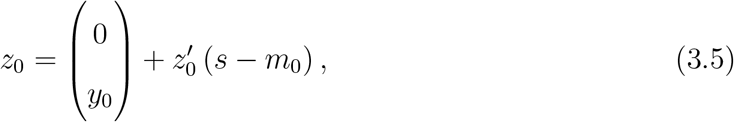

where the leading-order motor relative position, *m*_0_, and leading-order vertical position of the intersection, *y*_0_, complete the parameterisation. The leading-order ansatz is illustrated in Figure 3.1.

To obtain the leading-order solution, we consider the 𝒪(1) problem

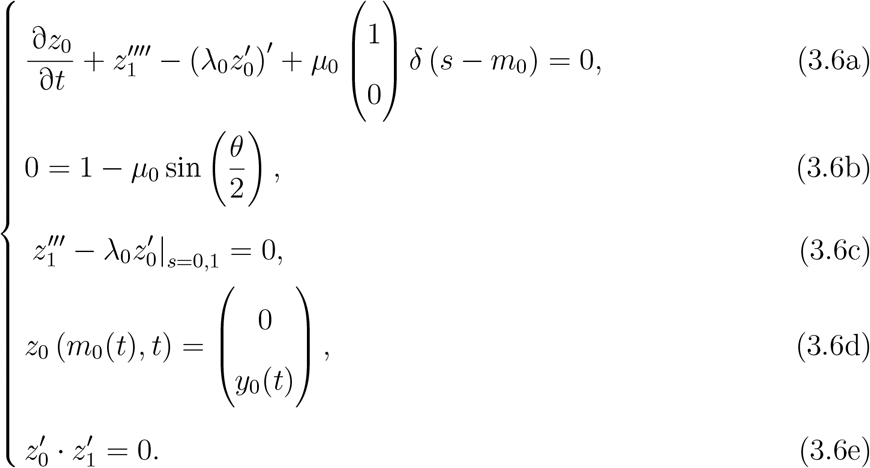

Equation (3.6a) and (3.6b) are the leading-order equations governing the filaments and motor respectively. Equation (3.6c) provides two boundary conditions, and the solution must also satisfy the ansatz (3.6d) and orthogonality constraint (3.6e). To proceed, we use the orthogonality condition (3.6e) to infer the ansatz

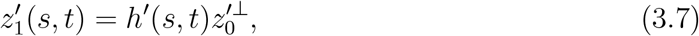

where *h*(*s, t*) is an arbitrary scalar function. Substituting the ansatzes (3.5) and (3.7) into the PDE for filament evolution (3.6a) enables us to solve for the leading-order quantities

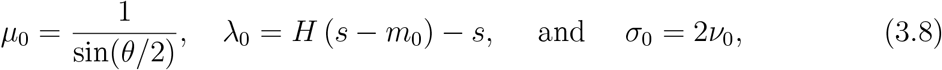

where *H* is the Heaviside step function, and *ν*_0_ = *m*_0_ − 1*/*2. Filament evolution then satisfies the ordinary differential equations

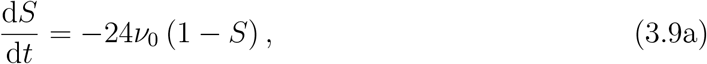

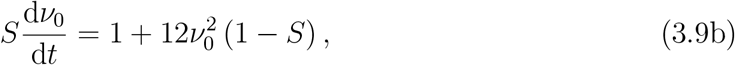

where *S* = sin^2^(*θ/*2). Since *z*_0_ is written in terms of the angle *θ* only, the system (3.9) determines *z*_0_. With *h*′ known, we subsequently obtain *z*_1_. Full details on this calculation are available in Appendix B.1.

An important property of the system (3.9) is that it is invariant under a change of variables that reverses the direction of time. If we introduce the reversed-time 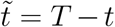 for an arbitrary constant *T*, we have 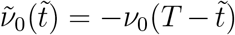 and 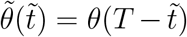, and subsequently 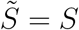. Consequently, if the motor is initially positioned at the pointed ends (*ν*_0_(0) = −1*/*2), and *T* denotes the time it reaches the barbed ends, then the time-aggregated stress vanishes,

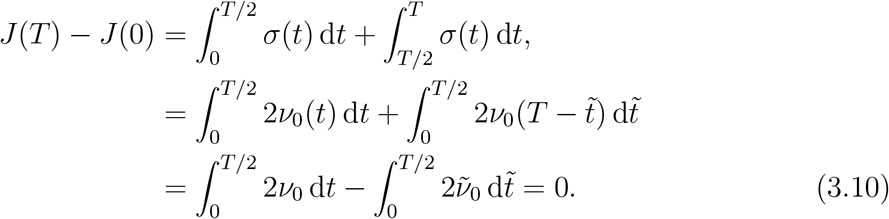

This is because the equations and initial conditions satisfied by *ν*_0_, *θ*, and 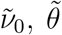 both coincide, and we have that 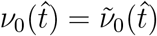 for all 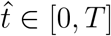 (see also numerical result shown in Figure 3.2c). This finding agrees with the previously reported results that rigid filaments with polarity-reversal symmetry produce zero net stress [25, 41].

**Figure 3.2:**
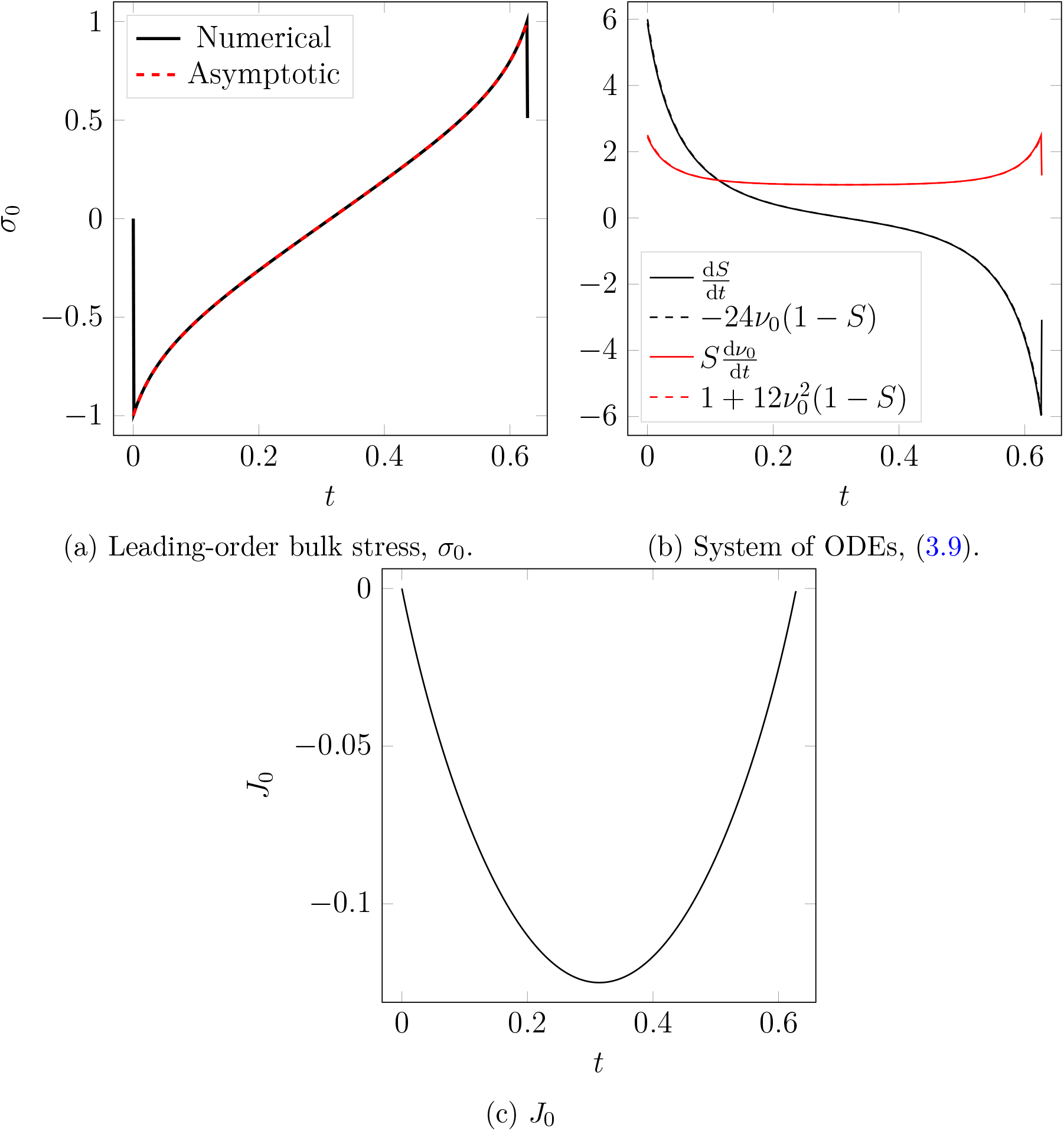
Comparison between a numerical solution with rigid filaments (*ε* = 1 × 10^−5^) and the leading-order asymptotic solution.

#### 3.1.2 First-Order Correction

The higher-order correction terms, *z*_1_, *σ*_1_ and *J*_1_, elucidate the effect of small, non-zero bending on filament evolution and stress. To obtain expressions for the first-order corrections to bulk stress and *J*, we substitute the asymptotic expansions (3.1) into the stress terms (2.30) and (2.31). This yields

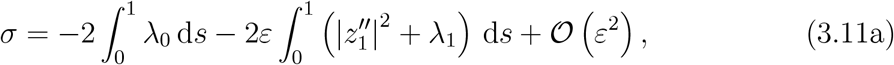

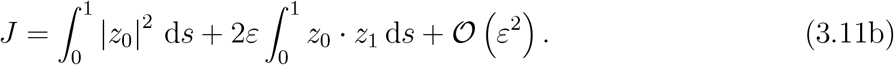

Matching coefficients of *ε* then yields

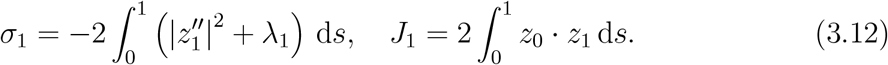

In addition, we use the PDE (3.6a) and the ansatz (3.7) to obtain an explicit expression for the curvature of *z*_1_,

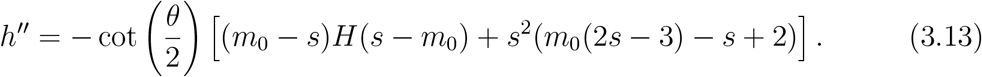

Since the first-order correction to stress, *σ*_1_, involves the currently unknown *λ*_1_, progress requires consideration of the 𝒪(*ε*) problem, which is

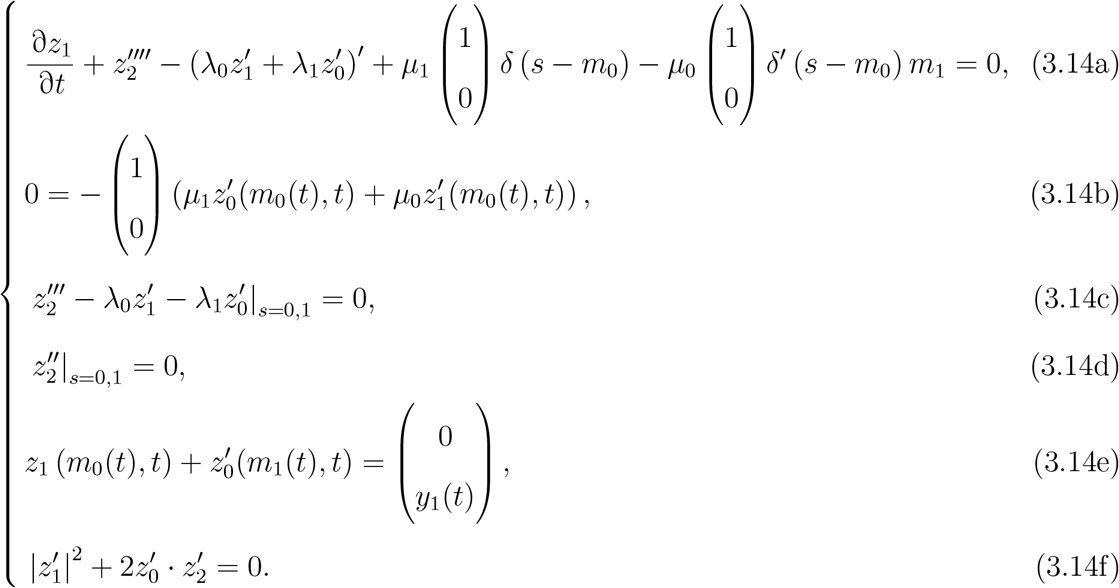

Obtaining the solution to (3.14) involves an intricate calculation based on the ansatz

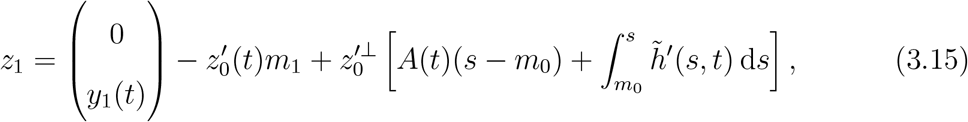

where *A*(*t*) is a (possibly time-dependent) constant of integration. The form of (3.15) arises from the ansatzes (3.7) and (3.14e), and gives rise to a system of equations for the degrees of freedom *A*(*t*), *y*_1_(*t*), and *m*_1_(*t*). We provide full details on the calculation to obtain this in Appendix B.2. A key result is the stress correction term,

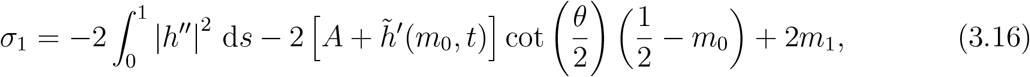

where 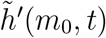 is given by

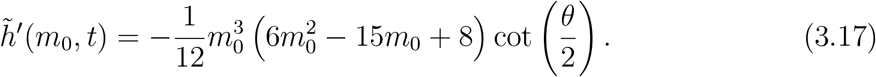

Similar to the system (3.9), we can obtain a system of differential equations to solve for *A*(*t*), *y*_1_(*t*), and *m*_1_(*t*). Since *h*″ and 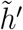 are in terms of the leading-order degrees of freedom *θ* and *m*_0_, we can subsequently compute *σ*_1_. However, the ODEs for *A*(*t*), *y*_1_(*t*), and *m*_1_(*t*) have no exact solution. Therefore, we continue our investigation using numerical solutions.

### 3.2 Numerical Solutions

We compute numerical solutions to the simplified governing equations for filament and motor positions (2.27) in Julia. Our numerical method involves minimising the time-discrete functional (2.13) with Δ*t* = 0.001. Energy minimisation is equivalent to a time-implicit numerical method for solving the dimensionless model (2.22). In our solutions, each filament has total length 1 μm [42], and consists of 50 equal-length line segments joined at nodes, about which segments can rotate. In Julia, we use the package `Optim.jl` [43] to obtain the minimiser using the limited-memory Broyden–Fletcher–Goldfarb–Shanno (LBFGS) method. After obtaining the minimiser, at each time step we use automatic differentiation (`ForwardDiff.jl`) of the functional (2.13) to compute the forces ***F***_*x*_ and ***F***_*y*_, and subsequently bulk stress *σ* (2.16).

#### 3.2.1 Comparison With Asymptotic Analysis

We begin by computing numerical solutions for two symmetric filaments with *m*(0) = 0, and *θ*(0) = *π/*2. Like the asymptotic analysis, we assume these filaments are initially rigid, *V*_*m*_ → ∞, and solve until the motor reaches the plus-end and detaches. First, we compute a solution for two rigid (*ε* = 1 × 10^−5^) filaments, to validate the leading-order bulk stress *σ*_0_ = 2*ν*_0_, and the solution to the system of ODEs (3.9), which governs *z*_0_. As Figures 3.2a and 3.2b show, for both of these we obtain agreement between the numerical solution and leading-order solution. Furthermore, Figure 3.2c illustrates the result from (3.10), namely that zero net stress is generated when a motor traverses two rigid filaments from the minus to plus-ends, *i*.*e. J*_0_(*T*) = *J*_0_(0) = 0, where *T* = 0.627 is the time at which the motor reaches the plus end.

Next, we solve the model with *ε* = 0.01 to validate the formulae for *h*″ and *σ*_1_, (3.13) and (3.16) respectively. The dynamics of the two filaments and motor are illustrated in Figure 3.3. As part of the solution, we compute *h*″ using the asymptotic formula (3.13) and numerical values of *θ* and *m*, and compare this with the numerical value for the curvature,

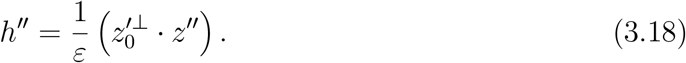

**Figure 3.3:**
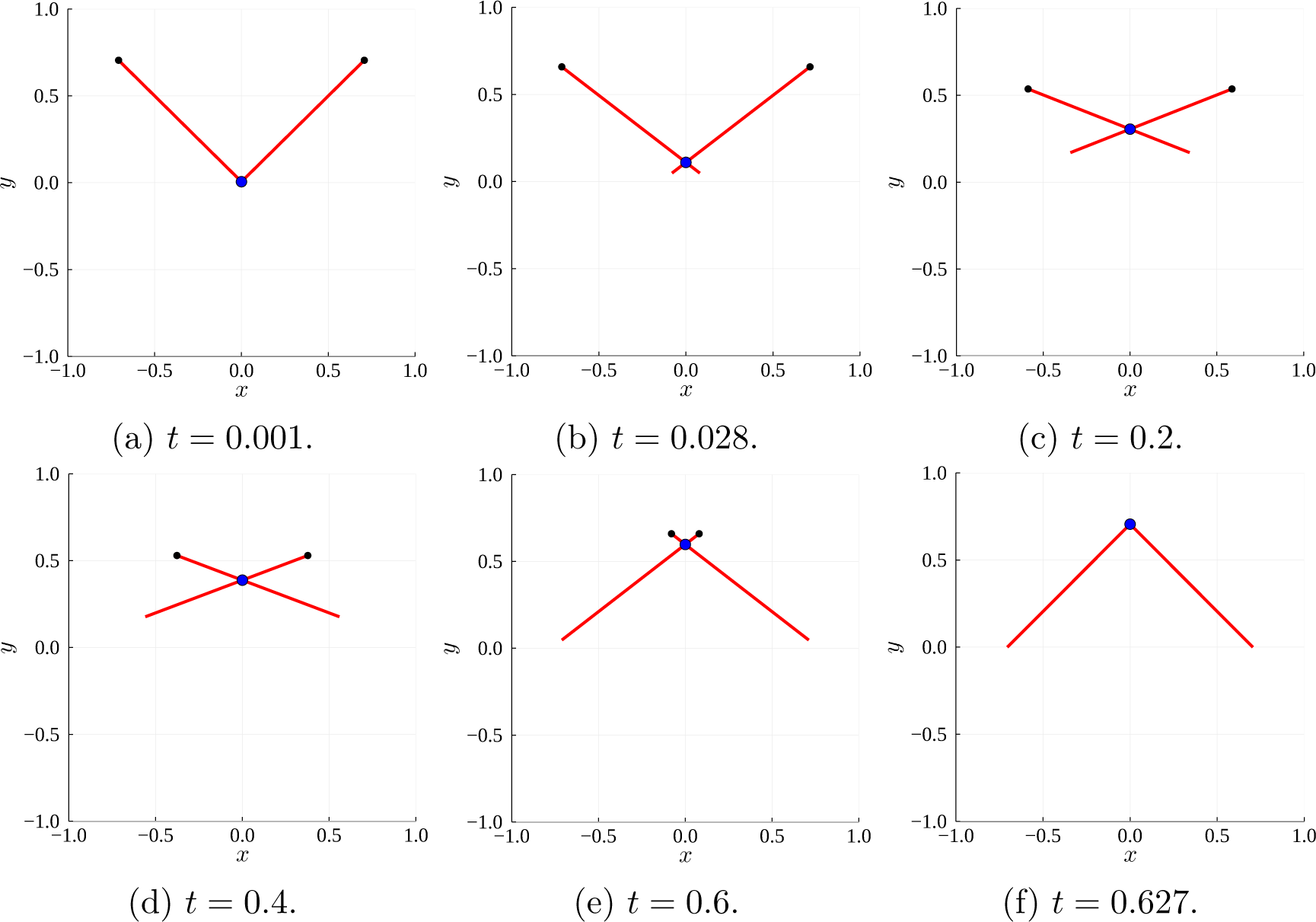
Numerical solution for the evolution of two actin filaments (red solid curves) with *ε* = 0.01. The black nodes indicate the filament plus ends, and the blue dot at the filament intersection represents the myosin motor.

As Figure 3.4 shows, we obtain agreement between the numerical and asymptotic results. The curvature formula (3.13) also reveals the shape that the two filaments adopt as they evolve (the qualitative pattern is easier to see in Figure 3.7). Initially, the filaments adopt a convex shape, as the positive curvature in Figure 3.4a shows. As the motor moves and pulls the filaments inwards, their shape changes to concave, as Figures 3.4c and 3.4d show. When the motor approaches the plus-end, the filaments return to a convex shape. The asymptotic result for *h*″ remains accurate for up to *ε* ∼ 𝒪(1), before breaking down for *ε* ∼ 𝒪(10).

**Figure 3.4:**
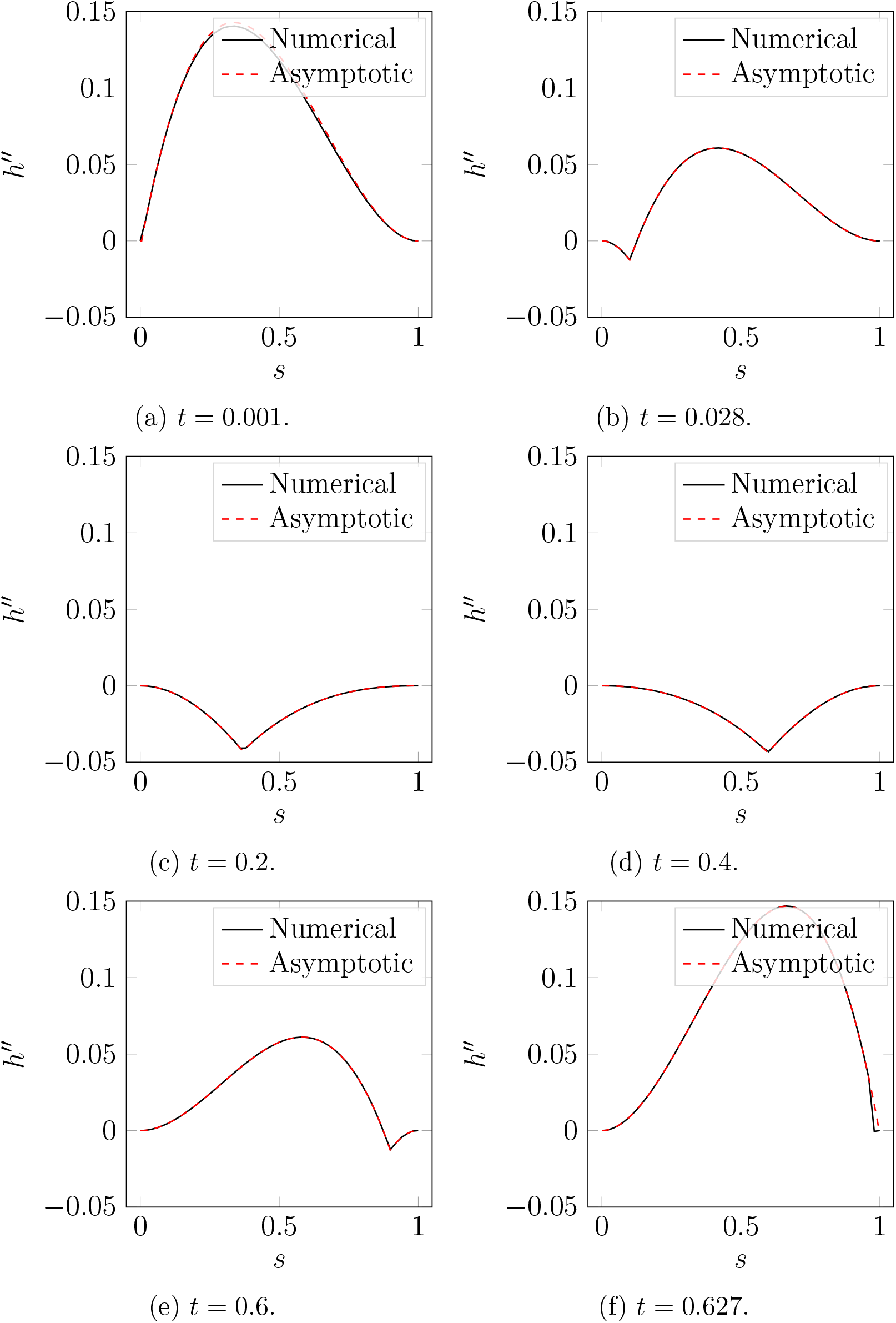
Numerical and asymptotic solutions for *h*″(*s, t*), the curvature of *z*_1_, in a numerical solution with *ε* = 0.01, and *θ*(0) = *π/*2.

We also use the numerical solution with *ε* = 0.01 to validate the formula for *σ*_1_, the first-order correction to bulk stress. At each time step, we compute the stress *σ*, and compare with the stress in a simulation with *ε* = 1 × 10^−4^, which we consider to be *σ*_0_ for rigid filaments. We then approximate the first-order correction as *σ*_1_ ≈ (*σ* − *σ*_0_)*/ε*, and present results in Figure 3.5a. For most values of *t*, it holds that *σ*_1_ *>* 0. In particular, larger positive values of *σ*_1_ occur close to *t* = 0 and *t* = *T*, or *m* = 0 and *m* = 1. Figure 3.5a is surprising, because it suggests the introduction of filament bending generates stresses that are biased to expansion. Similarly, as Figure 3.5b shows, the quantity *J*_1_(*T*) − *J*_1_(0) *>* 0, also suggesting net expansive bias. Based on this, one might conclude that bending cannot facilitate microscopic-scale contraction. However, we have not yet accounted for the changes in filament geometry, and how they influence motor dynamics. Further simulations in §3.2.2 will reveal this more clearly, and confirm that bending does facilitate net microscopic-scale contraction.

**Figure 3.5:**
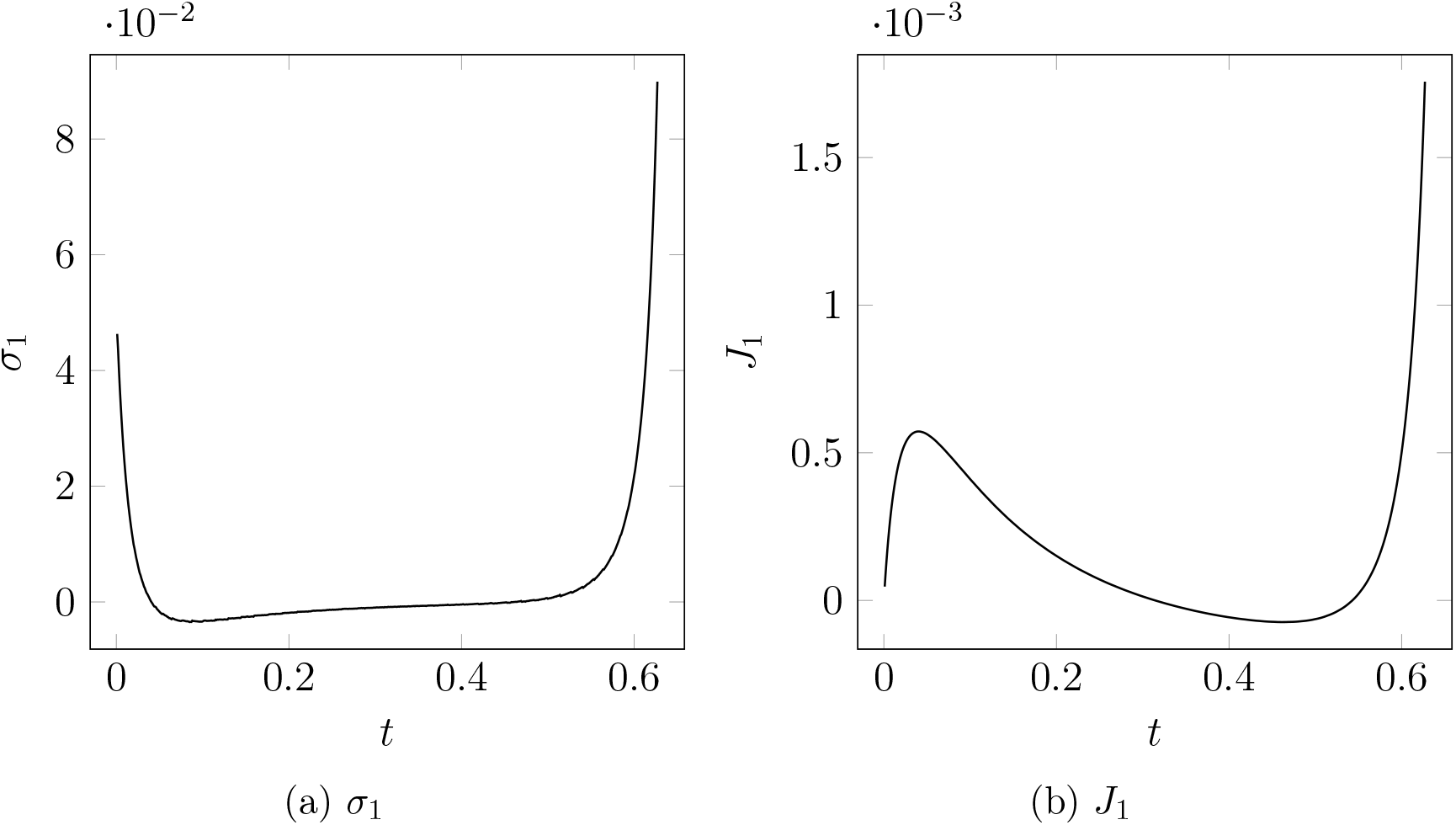
Calculation of *σ*_1_ ≈ (*σ* − *σ*_0_)*/ε* and *J*_1_ ≈ (*J* − *J*_0_)*/ε* in the numerical solution with *ε* = 0.01. The values of *σ*_0_ and *J*_0_ were obtained using a solution with *ε* = 1 × 10^−4^.

#### 3.2.2 Flexible Filament Solutions

We now consider numerical solutions beyond the *ε* ≪ 1 regime considered in the asymptotic analysis. These solutions are with the same conditions as Figure 3.3, where the motor is initially at the minus-ends of two symmetric filaments. We then solve the model until the motor reaches the plus-ends. Results are presented in Figure 3.6. The quantity *J* (*t*) measures the effect of *ε* on net stress, and is plotted in Figure 3.6a. For rigid filaments, we showed that *J* (*T*) − *J* (0) = 0, indicating zero net stress as the motor moved from the minus to the plus-ends. Since *J* (*T*) decreases as *ε* increases, the introduction of filament bending facilitates bias to contraction. This contractile bias is despite the quantities *σ*_1_(*T*) and *J*_1_(*T*) being positive, as in Figure 3.5a and 3.5b. Indeed, Figure 3.6b confirms that *σ*_1_ *>* 0, with stress increasing with *ε* close to *t* = 0 and *t* = *T*.

**Figure 3.6:**
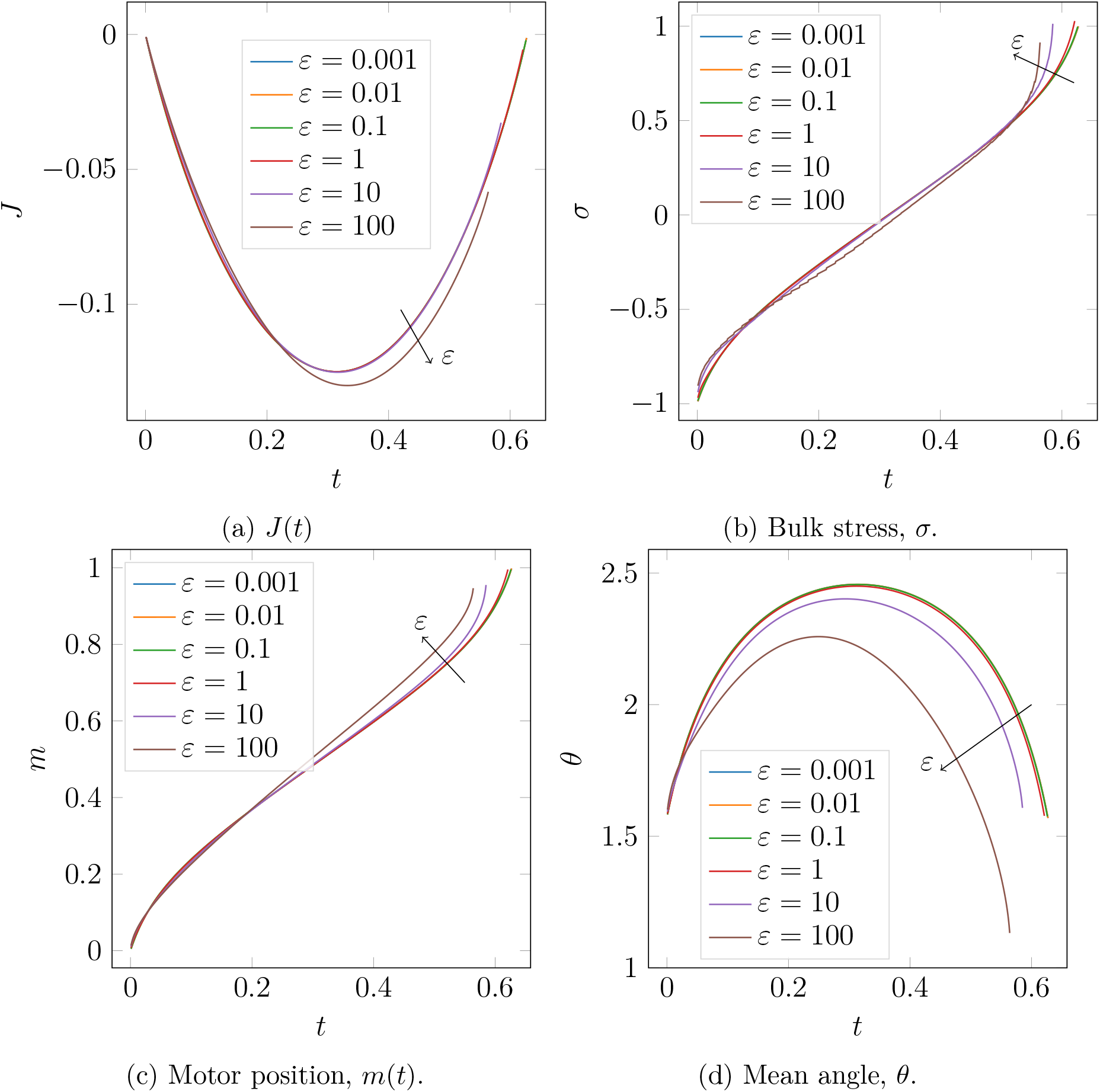
The effect of *ε* on quantities in solutions of two symmetric filaments. Solutions are computed with *m*(0) = 0, *θ*(0) = *π/*2, and proceed for all *t* ∈ [0, *T*(*ε*)] such that *m*(*t*) *<* 1. After this time *T*, the motor reaches the plus-end and detaches. Results are plotted for six values of *ε*, and arrows indicate the direction of increasing *ε*.

Semi-flexible filaments facilitate net contraction because bending breaks the polarity-reversal symmetry, and the resulting geometry favours contraction. As Figure 3.6c shows, with increasing *ε*, the myosin motor moves faster along the filaments and detaches earlier. The increase in motor speed is largest as the motor approaches the plus-ends, which Figure 3.6b shows is associated with expansion. As the motor approaches the plus-ends, the semi-flexible filaments adopt a convex shape that brings them closer to parallel at their tips, as illustrated in Figure 3.6d. This decreases the spring force through the motor, enabling it to move faster. Since the motor moves faster close to the plus-ends, the expansive component persists for shorter time than the contractile component. Consequently, the time-integrated stress *J* (*T*) − *J* (0) decreases as *ε* increases.

The results in Figure 3.6 are relevant for *in vivo* actin filaments, for which the parameters [42, 44–47] estimated in Tam, Mogilner, and Oelz [7] give *ε* = 68.5. To further our analysis, we compute a numerical solution with *ε* = 68.5 and *V*_*m*_ = 1, to investigate whether contraction persists after relaxing the assumption of infinite motor velocity. The evolution of these filaments is shown in Figure 3.7. Despite the slower motor speed, the evolution qualitatively follows the prediction from Figure 3.4. Filaments are initially convex, then become concave, and adopt a convex shape again as the motor approaches the plus-ends. As Figure 3.7f shows, the two filaments are curved when the motor reaches the plus-ends and detaches. To rule out the possibility that relaxation to straight configuration produces expansion that cancels out net contraction, we continued the simulation after motor detachment, until the filaments were again straight. We plot *σ*(*t*) and *J* (*t*) in Figure 3.8. Although relaxation to the straight configuration (shown in Figure 3.8a) generates a small amount of expansive stress, Figure 3.8c shows *J* (2) − *J* (0) *<* 0, suggesting net contraction. Thus, our proposed geometric mechanism for contraction remains relevant for realistic filament flexural rigidity and motor speed. Since actomyosin networks (for example those in the cortex) consist of many cross-linked two-filament assemblies, our mechanism provides a possible explanation for the microscopic origin of network-scale actomyosin contraction.

**Figure 3.7:**
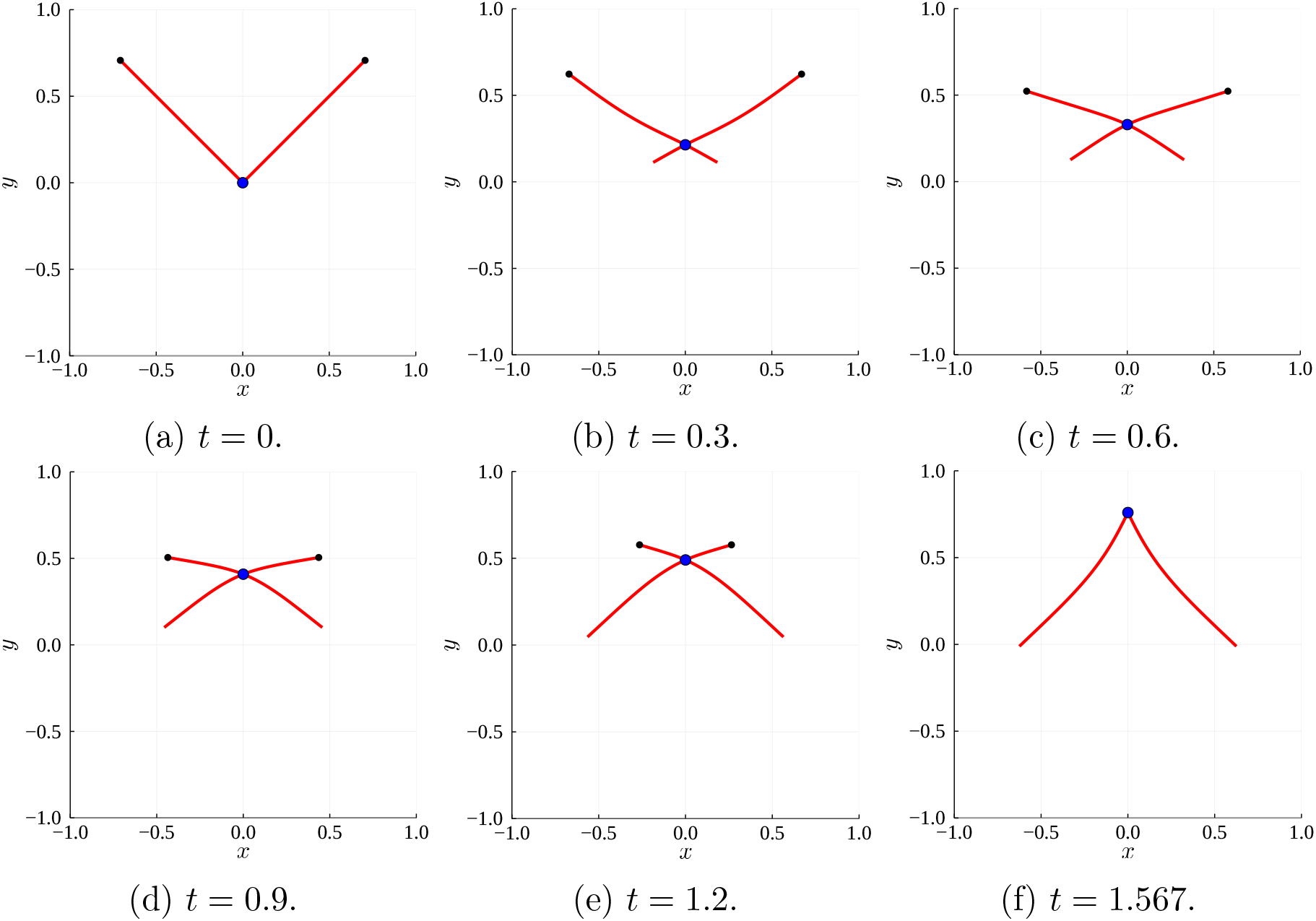
Numerical solution for the evolution of two flexible actin filaments (red solid curves) with *ε* = 68.5 and *V*_*m*_ = 1. The black nodes indicate the filament plus ends, and the blue dot at the filament intersection represents the myosin motor.

**Figure 3.8:**
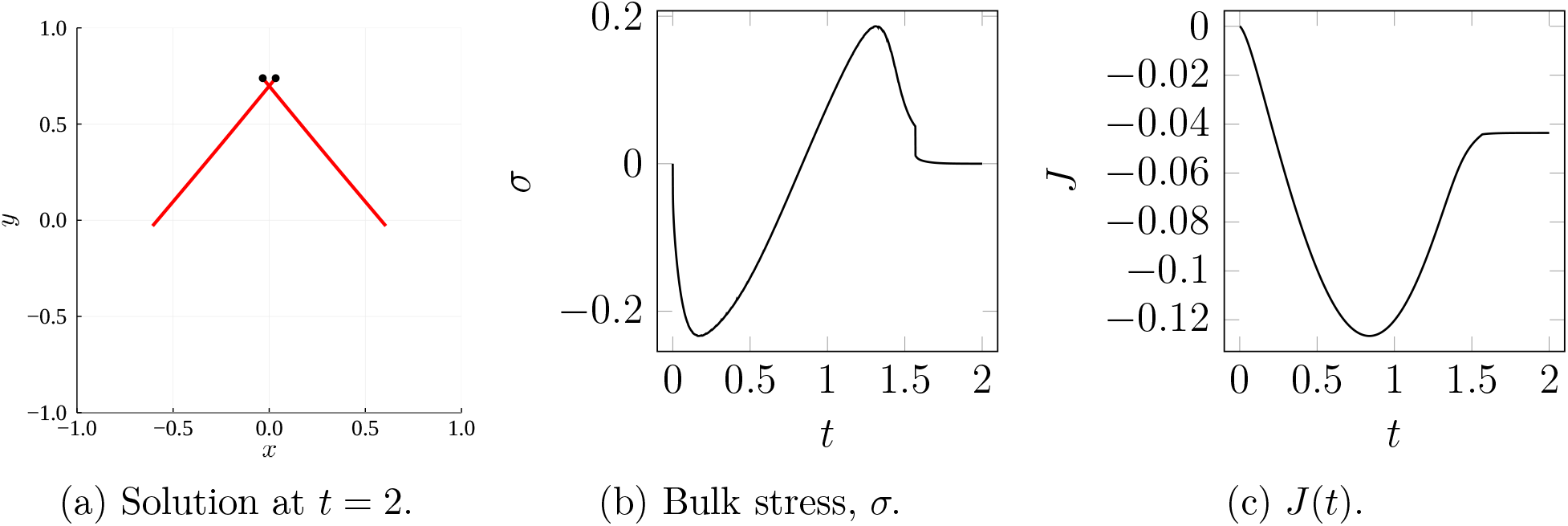
The final filament configuration, bulk stress, *σ*, and *J*(*t*) for the flexible filament (*ε* = 68.5 and *V*_*m*_ = 1) solution in Figure 3.7.

## 4 Conclusion

Understanding the origins of actomyosin contraction is an open problem in cellular biophysics, with implications for cell movement and division. In this paper, we presented a detailed investigation of how a two-filament-motor system generates microscopic contraction if the filaments are flexible. We first derived a partial differential equation model, and described a method of computing in-plane stress. We then applied asymptotic analysis to a symmetric system with infinite free-moving motor velocity. The leading-order solution showed that two rigid filaments do not generate net stress if the motor traverses the entire length of the filaments. However, the introduction of filament bending enables the two-filament–motor structure to generate net contraction. This is because bending breaks the polarity-reversal symmetry of rigid filaments. The resulting geometric asymmetry draws the plus-ends closer to parallel as the motor approaches. This facilitates faster motor movement when motors are close to filament plus-ends, and inhibits production of expansive stress.

Our analysis confirms that the microscopic dynamics of symmetric filament pairs and motors can explain contraction. We expect that the same mechanism also favours contraction in non-symmetric filament–motor assemblies and that, consequently, macroscopic contraction in disordered networks could arise from the accumulation of multiple filament pairs, without the need for nonlinear amplification of contractile stress. Nevertheless, the question of how these results apply to disordered networks remains open. In disordered networks, filament pairs cross at arbitrary angle and position, and interact with an active background of other filaments, rather than the passive medium considered in this work. Tam, Mogilner, and Oelz [7] confirmed that disordered networks described by the biomechanical model for semi-flexible filaments and motors presented in this study do contract. Another potential approach modelling disordered network contraction is to derive a coarse-grained, continuum model based on the assumption of infinite filament length [48]. This, and investigating how microscopic mechanics give rise to structures including stress fibres [49] and the contractile ring [42, 50], will be subjects of future work.

## Acknowledgements

The authors acknowledge funding from the Australian Research Council (ARC) Discovery Program (grant number DP180102956), awarded to D. B. O. and A. M.

## Declaration

The authors have no conflicts of interest to declare.

## A Mathematical Model Derivation

In this Appendix, we present a detailed derivation of the PDE model (2.10)–(2.12). We derive the system of force-balance PDEs using the variational principle. The functional for the structure consisting of a myosin motor attached to two semi-flexible actin filaments is

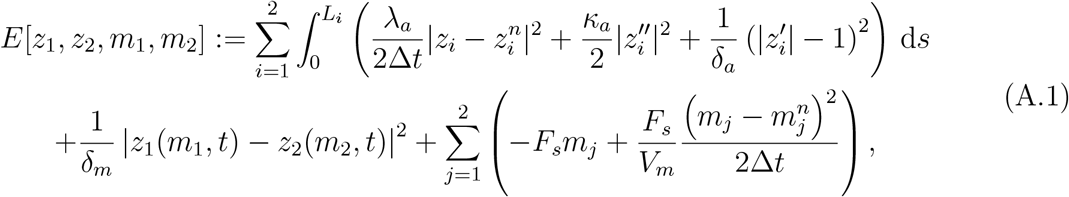

where *z*_*i*_(*s, t*) = (*x*_*i*_(*s, t*), *y*_*i*_(*s, t*)), for *i* = 1, 2, are the filament shapes and positions, *m*_1_(*t*), and *m*_2_(*t*) are the motor relative positions. According to the variational principle underlying the derivation, the solution at the next time step is such that the functional derivative with respect to all degrees of freedom vanishes, that is

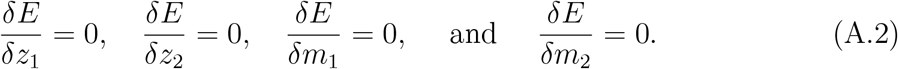

Evaluating each of (A.2) gives rise to the variational equations

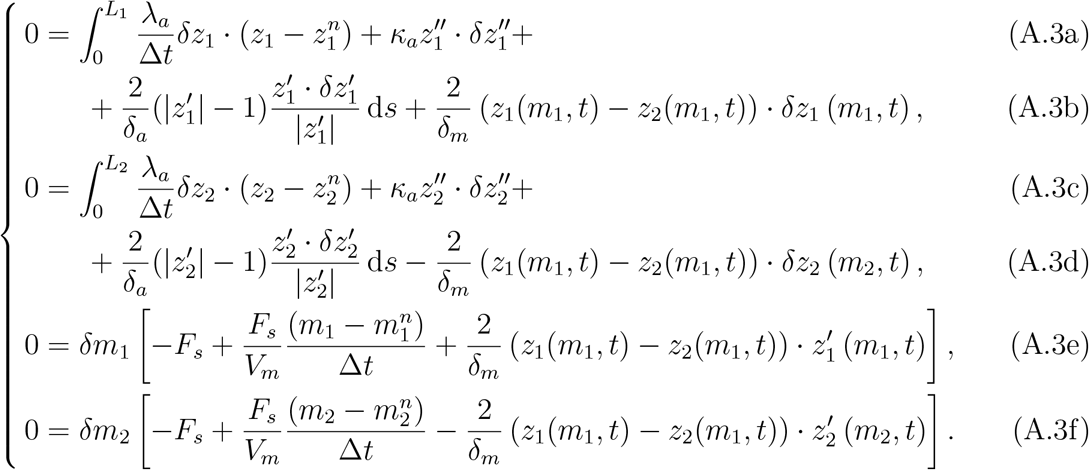

For the motor evolution equations (A.3e) and (A.3f), we can immediately apply the continuum limit Δ*t* → 0, for which 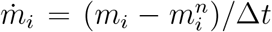 for *i* = 1, 2. In addition, we consider the limit *δ*_*m*_ → 0 and replace the forces due to motor stretching by the force *μ* which represents the Lagrange multiplier for the constraint

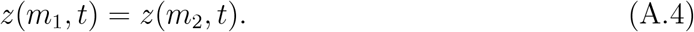

This yields the ordinary differential equations

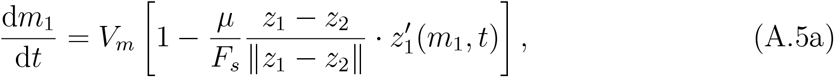

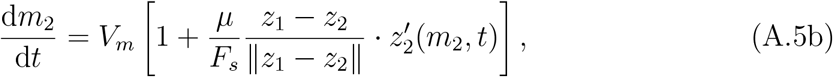

which are force-balance equations for the myosin motor positions *m*_1_(*t*) and *m*_2_(*t*). The equations (A.5) represent that unloaded motors (for which *μ* = 0) move at the free-moving velocity, *V*_*m*_. As the motor moves, it is exposed to stretching forces, with magnitude given by the Lagrange multiplier *μ*. The term involving the dot product is the projection of this force onto the direction of motor movement along the *i*-th filament. Assuming a linear force–velocity relation, the ratio of the term involving *μ* and the dot product to the stall force, *F*_*s*_, determines the reduction of motor speed due to stretching forces.

For the filament equations (A.3b) and (A.3d), in addition to *δ*_*m*_ → 0, we also let *δ*_*a*_ → 0 enforcing the constraints

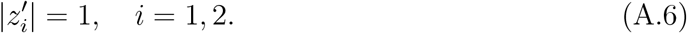

We write the limits of 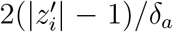 as *λ*_*i*_ and apply integration by parts to remove derivatives of *δz*_*i*_ from under the integral sign. This yields

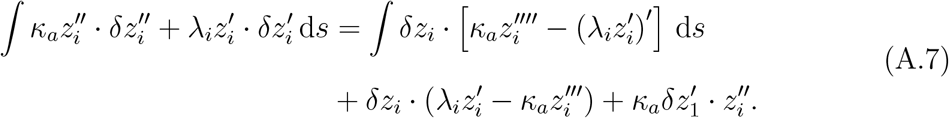

We then rewrite equations (A.3b) and (A.3d) as

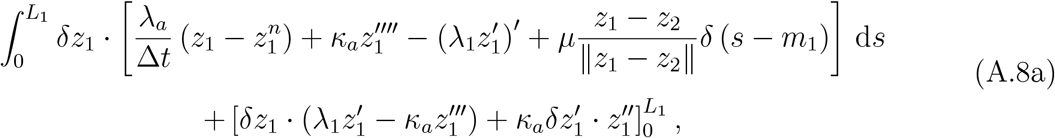

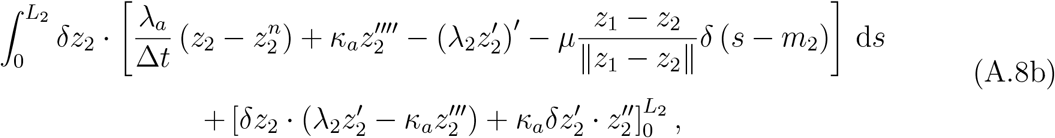

where *δ* is the Dirac delta function. The equations (A.8) enable us to derive the continuum governing equations and boundary conditions. First, we require the filaments to have zero curvature at their tips,

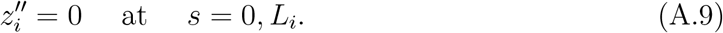

The remaining boundary terms in (A.8) then give rise to the conditions

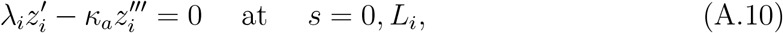

which specifies that the boundary values vanish at *s* = 0, *L*_*i*_. Next, we apply the fundamental lemma of the calculus of variations to the remaining integrals. In the continuum limit Δ*t* → 0 for which 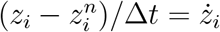, we obtain

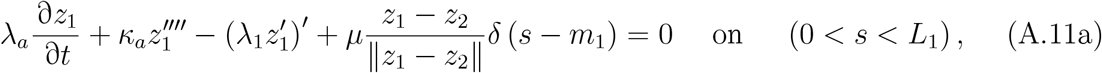

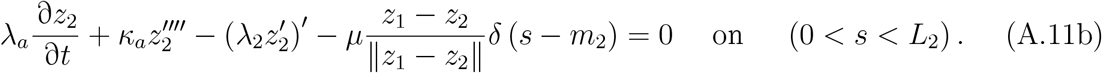

The differential equations (A.5) and (A.11), and conditions (A.9) and (A.10) as well as the constraints (A.4) and (A.6) then define a system of force-balance equations and boundary conditions that govern the evolution of two inextensible, semi-flexible filaments connected to an inextensible motor. This is the dimensional PDE model (2.10)–(2.12) given in the main text.

## B Asymptotic Analysis

In this Appendix, we present the asymptotic analysis of §3.1 in more detail. We first outline the method used to solve the leading-order problem (3.6), and subsequently consider the 𝒪(*ε*) problem (3.14) for the first-order corrections.

### B.1 Leading-Order Problem

We commence the analysis by considering the ansatz for rigid filaments. Taking the time derivative of (3.3) gives 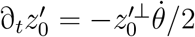, and taking four spatial derivatives of (3.7) yields 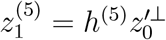. Next, we can differentiate the governing equation of 𝒪(1) (3.6a) once with respect to *s*, and substitute the two above expressions to obtain

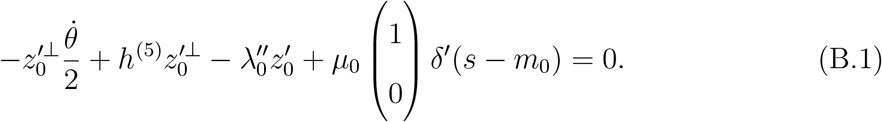

We can now multiply the boundary condition (3.6c) by 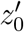, and use the property 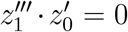, which follows from the orthogonality condition (3.6e), to obtain *λ*_0_(0) = *λ*_0_(1) = 0. Finally, multiplying (B.1) by 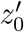 implies that

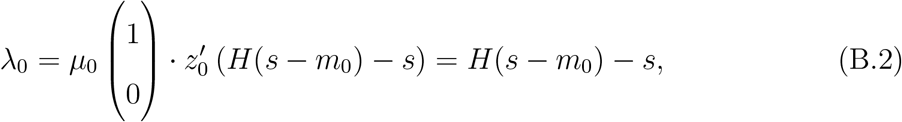

where *H*(*s*) denotes the Heaviside step function. The Lagrange multiplier *μ*_0_ = 1*/* sin(*θ/*2), by rearranging (3.6b). The leading-order bulk stress is then

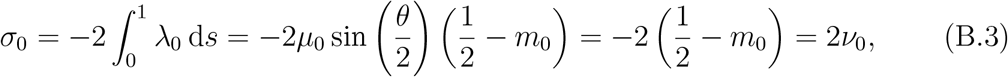

where *ν*_0_ = *m*_0_ − 1*/*2. This completes the derivation of the quantities listed in (3.8).

We now derive the ordinary differential equations (3.9) for *y*_0_, *θ*, and *ν*_0_, the three degrees of freedom that govern the leading-order filament position, *z*_0_. On taking the time derivative and variation, the ansatz (3.5) implies that

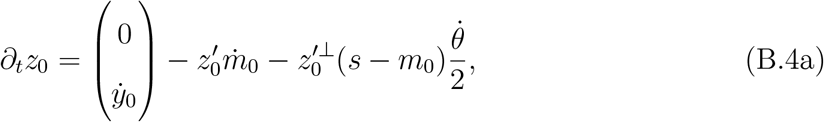

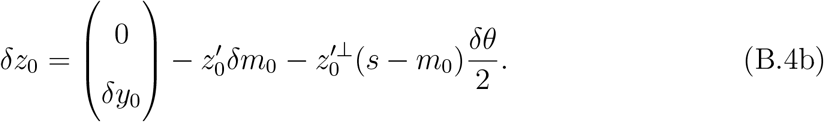

Integrating (3.6a) against *δz*_0_ then yields

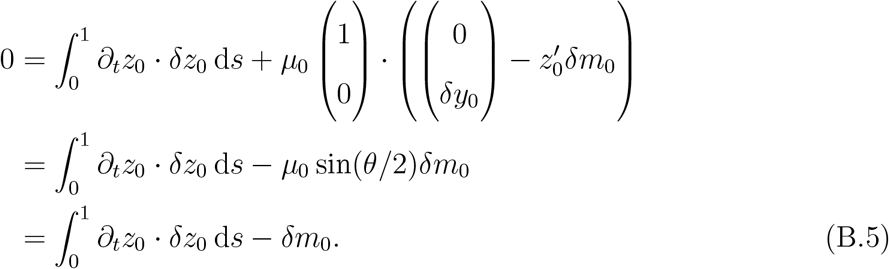

Substituting (B.4) into (B.5) then gives

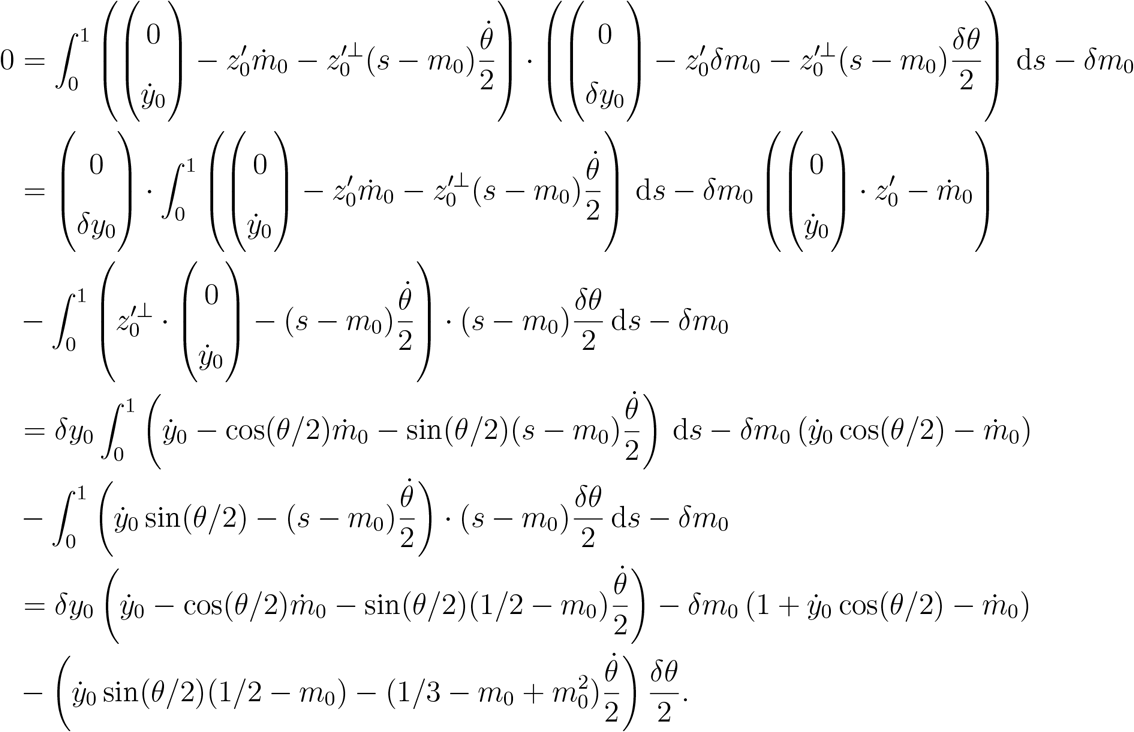

Collecting the coefficients of *δy*_0_, *δθ* and *δm*_0_, we obtain the system of differential equations (writing *ν*_0_ = *m*_0_ − 1*/*2)

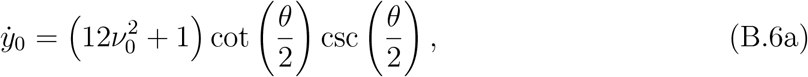

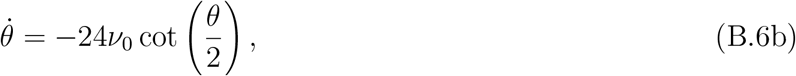

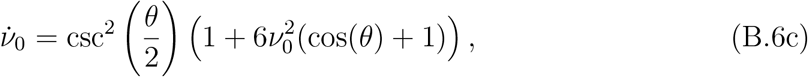

where csc(*ϕ*) = 1*/* sin(*ϕ*). The equations (B.6b) and (B.6c) for *θ* and *ν*_0_ are also independent of *y*_0_, suggesting that the solution is invariant to vertical translations. Furthermore, the trigonometric functions can be eliminated by writing *S* = sin^2^(*θ/*2), which yields

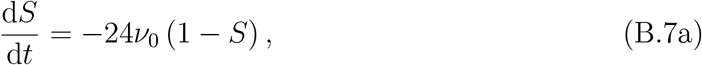

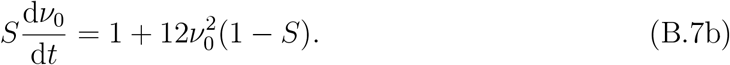

This completes the derivation of equation (3.9).

### B.2 First-Order Problem

To obtain the higher-order corrections, we first use the leading-order equation (B.1) and the ansatz (3.7) to solve for *h*, the curvature of *z*_1_. Multiplying (B.1) by 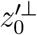 and using 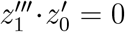 (which follows from the orthogonality condition (3.6e)), we obtain *h*‴(0) = *h*‴(1) = 0, and subsequently

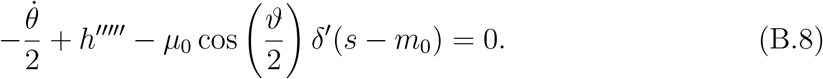

The boundary conditions *h*‴(0) = = *h*‴(1) imply that

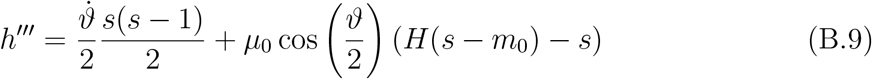

and furthermore (since 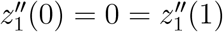), substituting for 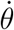 and *μ*_0_,

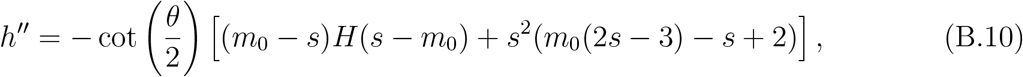

which is an expression for filament curvature, *h*″(*s, t*).

We now obtain the perturbation solution for the bulk stress, *σ*_1_, which requires knowledge of the quantities *μ*_1_ and *λ*_1_. First, we integrate (B.10) once with respect to *s*. This introduces another constant of integration, denoted *A*(*t*), which might be time-dependent and cannot be determined from boundary data. Consequently, we write

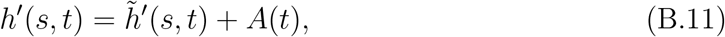

where

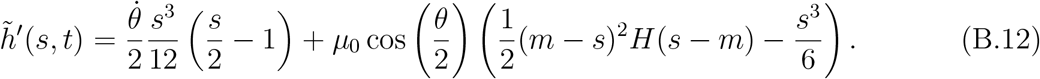

Collecting the coefficients of *ε* in the governing equations with asymptotic expansions, we obtain the 𝒪(*ε*) problem (3.14). On taking a derivative of (3.14a) and substituting 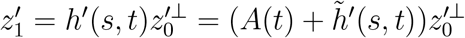, we obtain

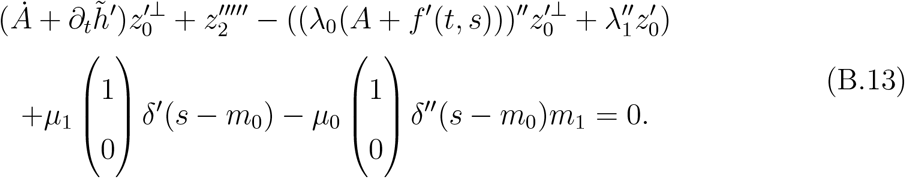

Expanding the inextensibility constraint (3.14f) implies that 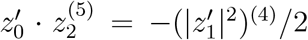. Multiplying (B.13) by 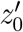, we obtain

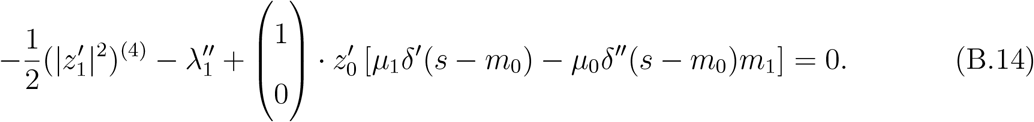

We can integrate (B.14) twice and apply the boundary conditions (3.14c) to determine the constants of integration. This yields

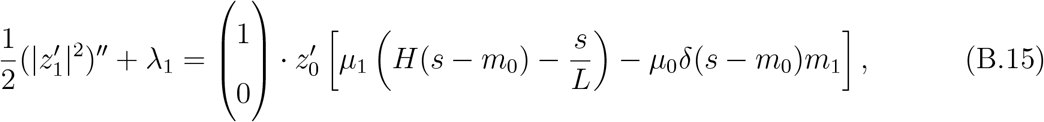

which we can rearrange to obtain *λ*_1_. To eliminate *μ*_1_ from (B.15), we use (3.14b) to infer an expression for *μ*_1_. Substituting the ansatzes (3.5) and (3.7) for *z*_0_ and *z*_1_ respectively, we obtain

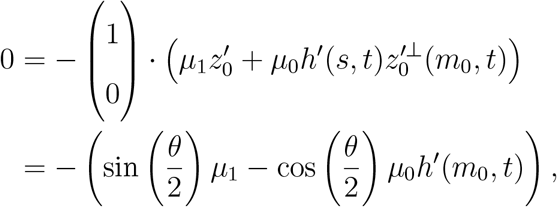

and therefore

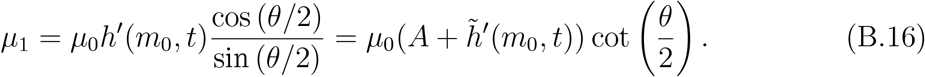

Substituting (B.16) into (B.15), we obtain

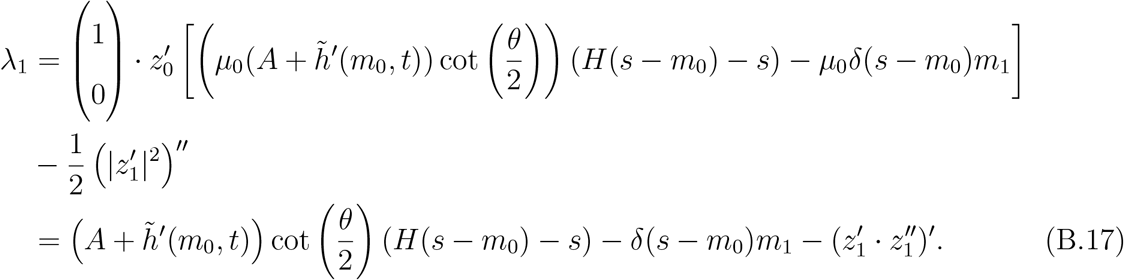

Using the simplified expression (B.17) for *λ*_1_, we can write the first-order perturbation of the bulk stress (3.12), given by

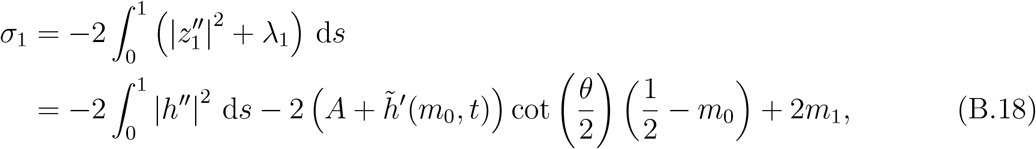

where 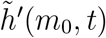 is

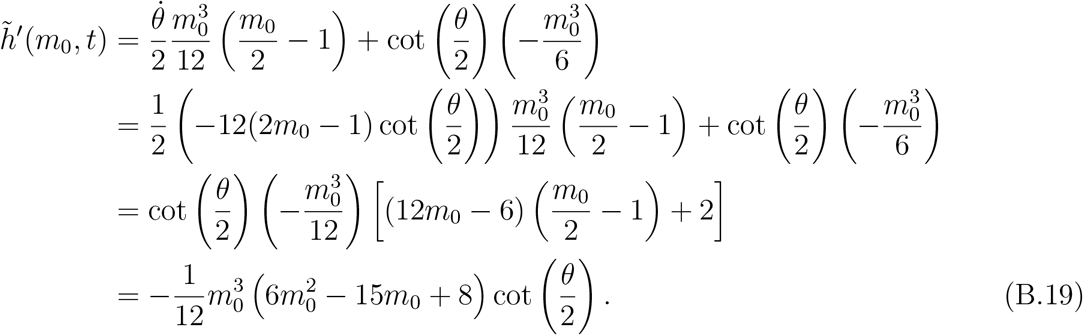

To solve for the first-order correction to the filament positions, *z*_1_, we require an initial condition, here denoted *z*_*I*,1_(*s*) = *z*_1_(*s*, 0). To determine the asymptotic expansion of the initial condition (2.28c), we return to the force-balance equations (2.1), and its equivalent time-discrete minimisation problem for the functional (2.2). In this expression, the drag component (2.3) dominates when Δ*t* is small. Therefore, we determine the leading order term in the asymptotic expansion of the initial condition *z*_*I*_ = *z*_*I*,0_ + *εz*_*I*,0_ + … as the best approximation of *z*_*I*_ in *L*^2^ among the straight fibres (3.5), that is

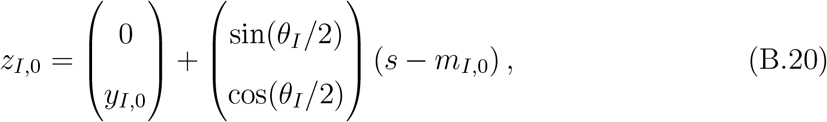

where

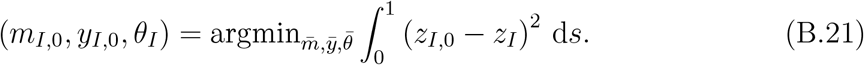

Since we focus on pairs of initially straight fibres in this study, we set *z*_*I*_ = *z*_*I*,0_.

A similar approach is available to determine *z*_*I*,1_. Using the ansatz (3.7), we have

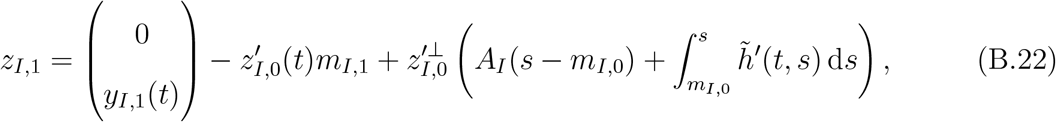

where

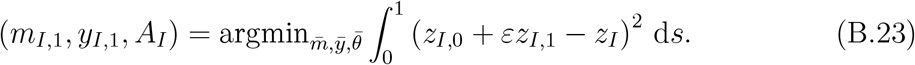

In the case where *z*_*I*,0_ = *z*_*I*_ the term *z*_*I*,1_ is minimal in *L*^2^. Then, the degrees of freedom *m*_*I*,1_, *y*_*I*,1_, and *A*_*I*_ can be computed using

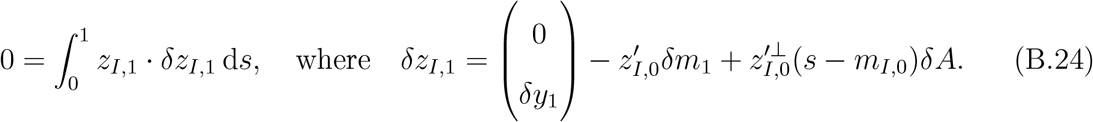

When we set 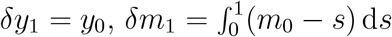, and *δA* = 0, we find that

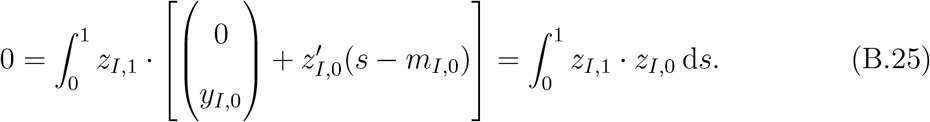

It therefore holds that *J*_1_(0) = 0.

To complete the derivation, we use the ansatz (3.7) with degrees of freedom *A*(*t*), *y*_1_(*t*), and *m*_1_(*t*). Its variation and time-derivative are given by

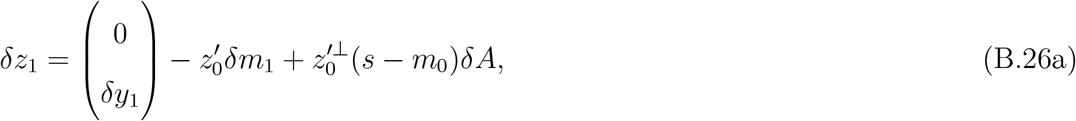

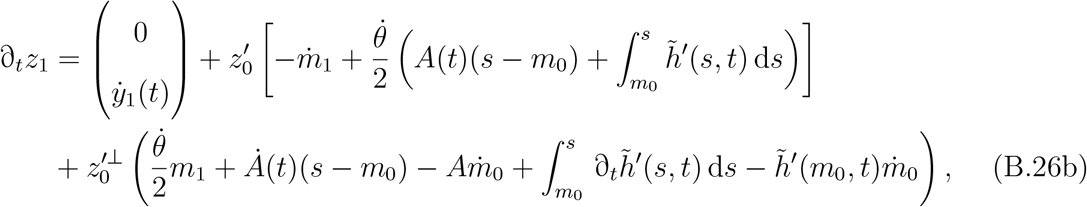

where 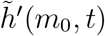 is given in (B.19). A system of differential equations for *m*_1_, *A* and *y*_1_ can then be found integrating (3.14a) against *δz*_1_. Using computer algebra, we obtained the system

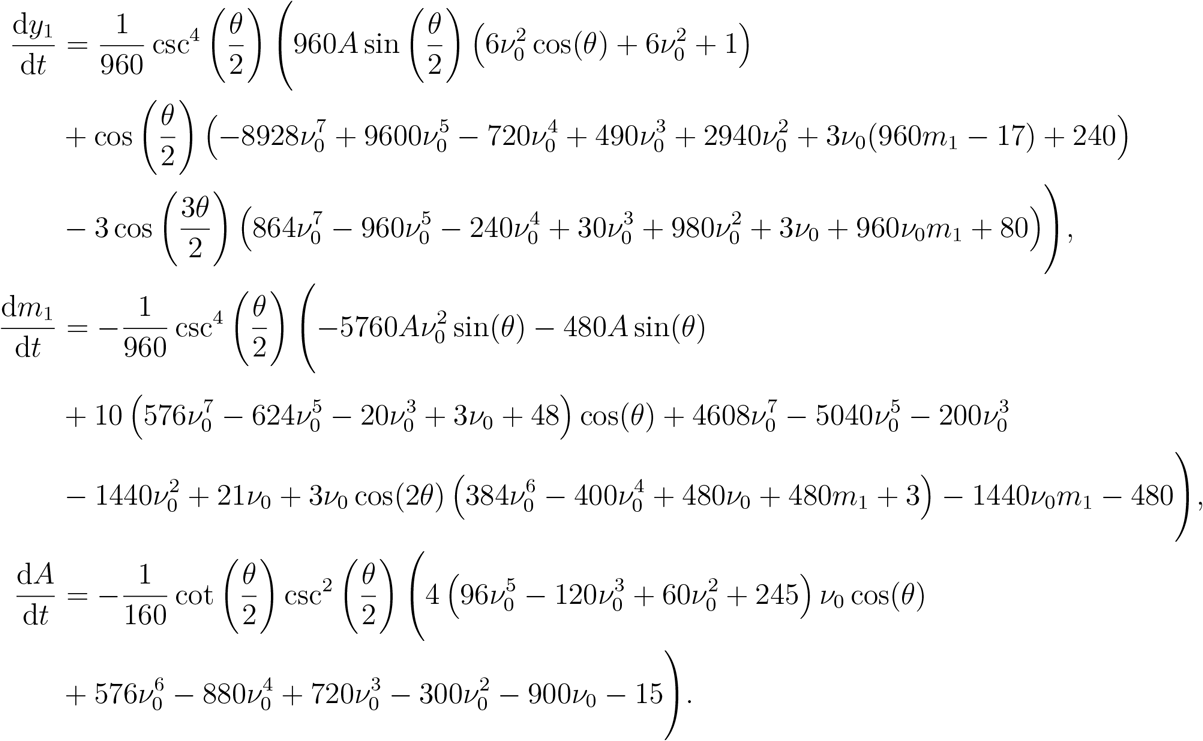

These ODEs govern the solution for *z*_1_, the first-order correction to the filament shape.

